# Characteristics of a SAR11 strain grown in batch and continuous culture

**DOI:** 10.1101/421644

**Authors:** Scott R. Grant, Matthew J. Church, Sara Ferrón, Edward A. Laws, Michael S. Rappé

## Abstract

In this study, a strain of SAR11 subgroup IIIa (termed HIMB114) isolated from the tropical Pacific Ocean was grown in seawater-based batch and continuous culture in order to quantify cellular features and metabolism relevant to SAR11 ecology. We report the first direct measurements of cellular elemental quotas for nitrogen (N) and phosphorus (P) for SAR11: 1.4 ± 0.9 fg N and 0.44 ± 0.01 fg P, respectively, that were consistent with the small size of HIMB114 cells (average volume of 0.09 µm^3^). However, the mean carbon (C) cellular quota of 50 ± 47 fg C was anomalously high, but variable. Rates of phosphate (PO_4_^3-^) uptake measured from both batch and continuous cultures were exceptionally slow: in chemostats growing at 0.3 d^−1^, HIMB114 took up 1.1 ± 0.3 amol P cell^−1^ d^−1^, suggesting that <30% of the cellular P requirement of HIMB114 was met by PO_4_^3-^ assimilation. The mean rate of leucine incorporation, a measure of bacterial production, during late log phase growth of batch HIMB114 cultures was 0.042 ± 0.02 amol Leu cell^−1^ h^−1^. While only weakly correlated with changes in specific growth rates, the onset of stationary phase resulted in decreases in cell-specific leucine incorporation that were proportional to changes in growth rate. Rates of cellular production, respiratory oxygen consumption, and changes in total organic C concentrations constrained cellular growth efficiencies to 13 ± 4%. Hence, despite the small, streamlined genome and diminutively sized cells, SAR11 strain HIMB114 appears to grow at efficiencies similar to naturally occurring bacterioplankton communities.

**Importance:** While SAR11 bacteria contribute a significant fraction to the total picoplankton biomass in the ocean and likely are major players in organic C and nutrient cycling, the cellular characteristics and metabolic features of most lineages have either been only hypothesized from genomes or otherwise not measured in controlled laboratory experimentation. The dearth of data on even the most basic characteristics for what is arguably the most abundant heterotroph in seawater has limited the specific consideration of SAR11 in ocean ecosystem modeling efforts. In this study, we provide measures of cellular P, N, C, aerobic respiration and bacterial production for a SAR11 strain growing in natural seawater media that can be used to directly relate these features of SAR11 to biogeochemical cycling in the oceans. Through the development of a chemostat system to measure nutrient uptake during steady-state growth, we have also documented inorganic P uptake rates that allude to the importance of organic phosphorous to meet cellular P demands, even in the presence of non-limiting PO_4_^3-^ concentrations.

## Introduction

The SAR11 bacterial lineage is a genetically diverse clade of aquatic, free-living cells with compact, streamlined genomes, found broadly distributed throughout the oceans (1). They are also among the smallest free-living cells from the ocean that have isolated strains available to study in the laboratory (2). Typical biovolumes for healthy SAR11 cells range from 0.015 to 0.058 µm^3^ (3), and possess a crescent-shaped morphology (2-5). Small cells are thought to have an advantage in oligotrophic environments where they should be able to out-compete larger osmotrophs for nutrients relative to their requirements for growth, ascribed to the importance of having a large surface area to volume ratio (6).

Culture studies examining the physiology of SAR11 strains have provided a number of unexpected discoveries and valuable insights into the metabolism of the clade (1). Directed by clues generated from genome analysis indicating that a number of metabolic pathways common to chemoheterotrophs were incomplete or missing, subsequent culture studies led to evidence of unusual growth requirements for SAR11 (e.g. 5, 7-12). For example, evidence of an incomplete assimilatory sulfate reduction pathway led Tripp and colleagues to the discovery that SAR11 strain HTCC1062 requires a source of reduced sulfur for growth, which could be satisfied by methionine or dimethylsulphoniopropionate (7). Further investigations showed that SAR11 had a variant of the standard glycolysis pathways, with nonconserved ability of SAR11 strains to oxidize simple sugars, while low molecular weight organic acids were shown to be important C sources for many SAR11 strains (9). In subsequent experiments, Carini and colleagues were able to successfully grow SAR11 strain HTCC1062 on a novel defined artificial seawater medium with pyruvate serving as a source of C, methionine as a sole sulfur source, glycine as a necessary amino acid, along with standard base salts, inorganic macro-nutrients PO_4_^3-^ and ammonium (NH_4_^+^), and micro-nutrient trace metal and vitamin additions (5). Laboratory experiments with isolated SAR11 strains have primarily focused on representatives from the SAR11 subclade Ia, which includes the majority of isolates including ‘*Candidatus* Pelagibacter ubique’ strain HTCC1062 (2), with little information from representatives of other SAR11 subclades.

Recent studies suggest that the type of P available, whether present as PO_4_^3-^ or dissolved organic P (DOP), is an important control on microbial niche partitioning in the sea (13, 14). The Global Ocean Sampling (GOS) expedition, an extensive metagenomic survey of marine surface waters, revealed that genes from the high-affinity PO_4_^3-^ transport system (*pstS*) most closely matching sequenced *Prochlorococcus* and SAR11 genes, were among the most highly recruited annotated genes (15). Moreover, during the GOS expedition, representation of *pstS* genes were the single most significant difference between the tropical Atlantic and equatorial Pacific samples, varying by a factor of more than seven in relative abundance (15). Studies of culture representatives of *Prochlorococcus*, the most abundant oxygenic photoautotroph in the ocean, confirm that there appear to be substantial differences in the presence, topology, and regulation of genes thought to be involved in P acquisition between strains of *Prochlorococcus* (16), with different strains able to metabolize inorganic vs. labile organic P compounds. Finally, in a gene content comparison of whole population genomes of *Prochlorococcus* and SAR11 between microbial communities inhabiting the well-known stations of the Hawaii Ocean Time-series (HOT) program (North Pacific) and the Bermuda Atlantic Time-series Study (BATS) (North Atlantic), Coleman and Chisholm found that of the 1.8% of gene clusters which had significant abundance differences between the Atlantic and Pacific populations, 87% of those genes were involved in PO_4_^3-^ or phosphonate metabolism (17).

Motivated by the intriguing evidence that P acquisition strategies are under strong selection pressure and may be a potential dimension over which SAR11 lineages are differentiated, this study sought to investigate the uptake capability of the SAR11 subclade IIIa isolate HIMB114 for PO_4_^3-^. Because SAR11 bacteria characteristically dominate marine planktonic microbial communities, it is also a notable deficiency that typical parameters needed to model their growth and response under variable environmental conditions are not yet available. Thus, this study also sought to measure a number of basic cellular properties such as elemental composition, and physiological rate measurements including cellular production, respiration, and growth efficiency. In the process, a continuous culture of an axenic SAR11 strain was developed for the first time, enabling assessment of many of these physiological features under defined growth conditions.

## Results

### Culture growth and cell size

In natural seawater-based growth media, strain HIMB114 reached a maximum specific growth rate of 1.2 d^−1^ and yielded 5-8 ×10^5^ cells mL^−1^ (Fig. S1). HIMB114 cells were observed to have a crescent-shaped morphology that was consistent with previous microscopic observations of SAR11 (Fig. 1). Elongated cells of HIMB114 were observed that consisted of spirillum morphologies of two to four “regular” (i.e. recently divided) single-cell lengths. These longer cell morphologies were a small fraction (few percent) of the cells during exponential growth phase, but became an increasing percentage (up to 30%) of cells as the culture entered into stationary phase. Distributions of cell size parameters for length, width, and biovolume were all non-normal and fit as log-normal distributions to calculate the most frequent and mean size parameter values (Fig. S2). The mean of the distribution was used when normalizing any quantities to a cell size parameter (length 1.07 µm; width 0.32 µm; volume 0.09 µm^3^).

**Figure 1.**
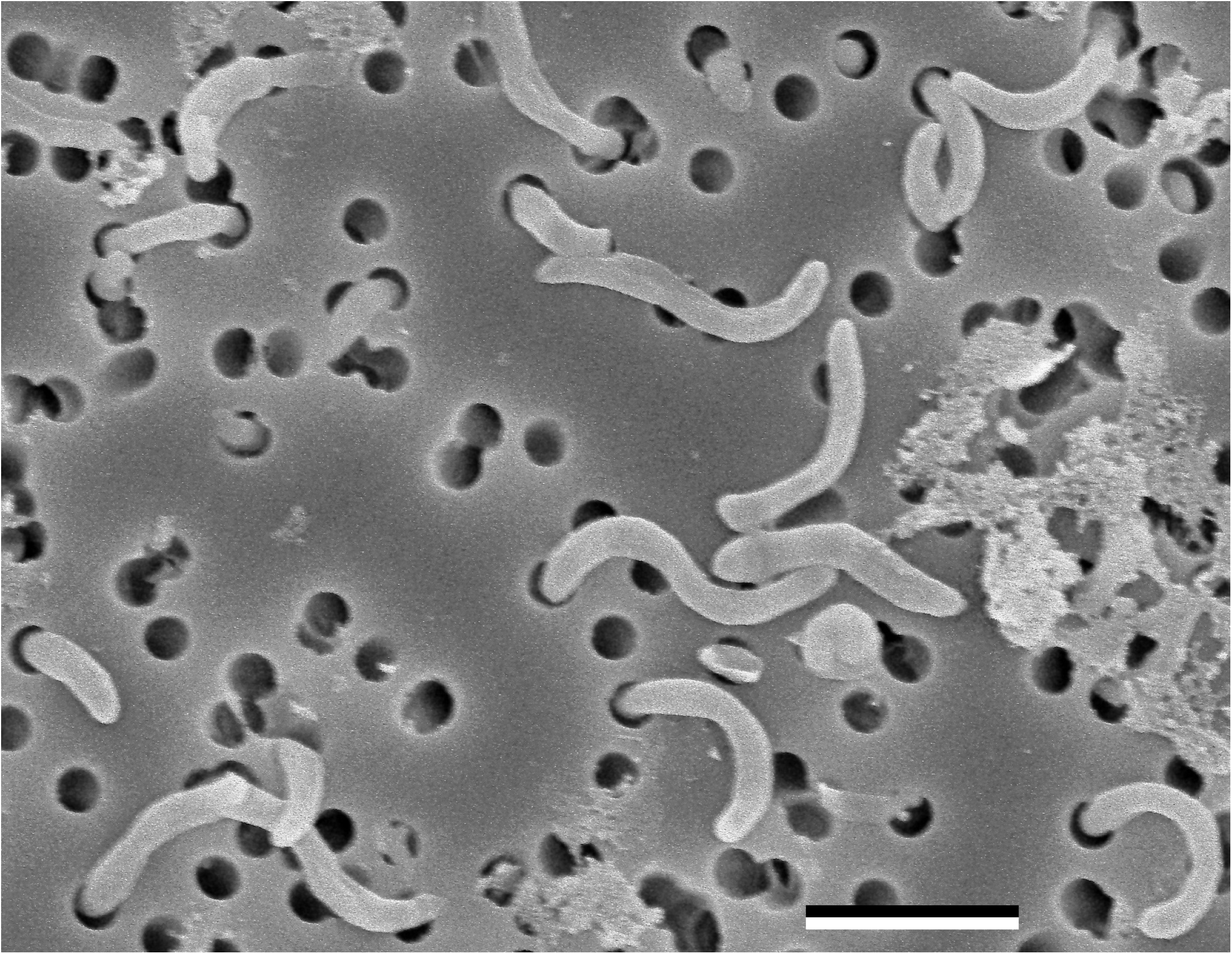
Scanning electron micrographs of HIMB114 cells growing in early stationary phase batch culture. The scale bar corresponds to 1 µm.

### Cellular elemental quotas

The P cell quota for HIMB114 measured for batch cultures in early stationary phase was 14.2 ± 0.4 amol P cell^−1^ (mean ± s.d.; n=6) or 0.44 fg P cell^−1^, with a mean precision of 2% for triplicate 4–5 L culture volumes. The particulate P controls made from spent media were 5% of that measured for the cellular biomass collected on their corresponding 0.2 µm pore-sized membrane filter. Hence, the modified method for measuring particulate P on 47 mm diameter PC membranes described in the Materials and Methods appeared to work well.

Filtered media blanks for particulate C and N were high relative to the sample signal, and increased with the volume of media filtered (Fig. S3). Because complete saturation was not conclusive even at a media blank volume of 10 L, rectangular hyperbolic saturation functions were fit by non-linear least squares regression to the blank C and N data versus filtered media volume in order to extrapolate the associated blank values for the 30 L of total volume filtered (Fig. S3). After normalizing to the total number of cells captured on each filter, the mean C cell quota was 50 ± 47 fg C cell^−1^ (mean ± s.d.; n=3) or 4.2 fmol C cell^−1^, and the mean N cell quota was 1.4 ± 0.9 fg N cell^−1^ (mean ± s.d.; n=3) or 0.1 fmol N cell^−1^.

### PO_4_^3-^ uptake in batch and continuous culture

Rates of PO_4_^3-^ uptake measured by ^33^P-radiotracer for a continuous culture of HIMB114 were extremely slow (Fig. 2A). Of the five PO_4_^3-^ uptake rate time course measurements performed from the chemostat cultures over 4 to 6 h at ambient (100 nmol L^−1^) phosphate concentrations, the mean specific uptake rate was 0.007 ± 0.0025 d^−1^ (mean ± s.d.; n=5), with a mean coefficient of determination of uptake vs. time of 0.97 (Fig. 2A). This corresponds to a mean PO_4_^3-^ turnover time (T_P_) of 160 ± 50 days (mean ± s.d.; n=5), or a bulk PO_4_^3-^ uptake rate of 0.68 nmol L^−1^ P d^−1^. In cell specific units, HIMB114 took up 1.1 ± 0.3 amol P cell^−1^ d^−1^ (mean ± s.d.; n=5), or less than 10% of the cellular P quota per day, despite growing at a rate of 0.3 d^−1^. Phosphate uptake kinetics measured for the chemostat culture averaged 0.4 ± 0.09 nmol L^−1^ P d^−1^ (mean ± s.d.; n=9) across all PO_4_^3-^ additions, showing no significant correlation between uptake rate and PO_4_^3-^ concentration within the error of the measurements (Fig. 2B). This observation most likely reflects the fact that the culture was not P-limited; an interpretation confirmed by the fact that PO_4_^3-^ additions to batch cultures entering stationary phase had no affect on growth (not shown).

**Figure 2.**
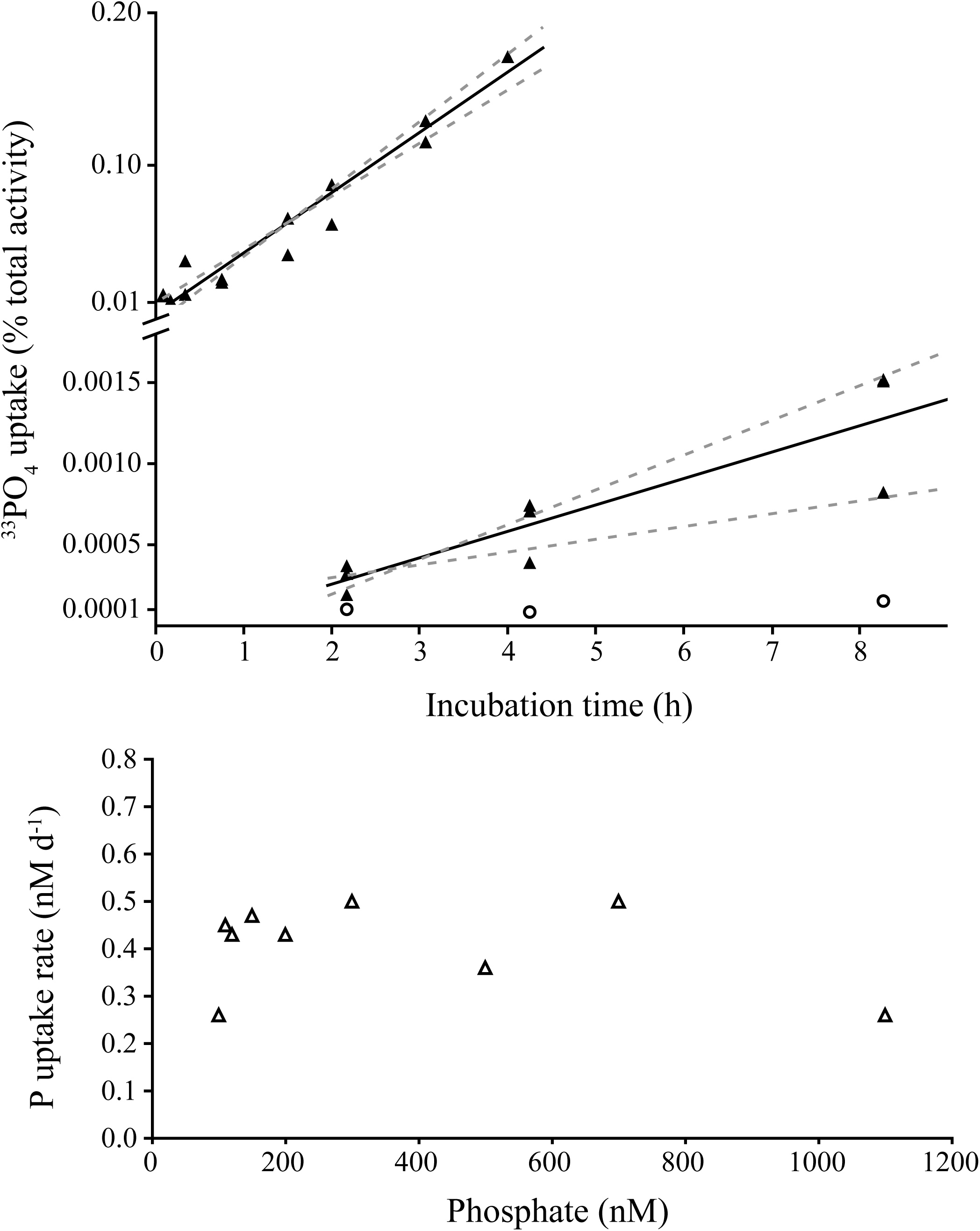
Rates of PO_4_^3-^ uptake by SAR11 strain HIMB114. (A) Time course measurements of ^33^P-phosphate uptake in a chemostat culture of HIMB114 (upper) as well as a batch culture (lower) with blank controls (circles). Solid lines indicate the linear least squares regression, while dashed lines indicate the 95% prediction confidence bands. (B) ^33^P-phosphate uptake kinetics for a chemostat culture of HIMB114, calculated from single time point, 22-hour incubations across a range of phosphate concentrations.

For HIMB114 grown under batch conditions, PO_4_^3-^ uptake rates were also extremely low (Fig. 2A). Measured during late exponential phase for a culture growing at 1.02 d^−1^, the highest specific uptake rate measured was 4×10^−5^ d^−1^, equivalent to a turnover time of the PO_4_^3-^ pool of 70 years (ranging 50 to 100 years). In bulk units, the maximum PO_4_^3-^ uptake rate for the batch cultures in late exponential growth was 6 pmol P L^−1^ d^−1^. To confirm that the cells were actively growing, leucine incorporation measurements were conducted at the same time as the PO_4_^3-^ uptake measurements (described in greater detail below). The resulting production rate was 37 ± 2.6 nmol C L^−1^ d^−1^ (mean ± s.d.; n=4) that, when converted to P units using a 50:1 C:P molar ratio, yields a P requirement of 0.7 nmol P L^−1^ d^−1^. Given that the measured bulk PO_4_^3-^ uptake rate was 6 pmol P L^−1^ d^−1^ (or ∼1% of the requirement), such results suggest PO_4_^3-^ was not the primary source of P for HIMB114 growing on natural seawater based media containing 100-150 nmol PO_4_ L^−1^.

### Chemostat steady-state theory

The theoretical expectations for PO_4_^3-^ uptake rate and turnover time measurements for the chemostat system are fairly well constrained, much better than for batch culture growth, because steady state theory may be applied (18). The cell specific uptake rate is the product of the specific growth rate (*µ*) with the cellular P quota (*Q*_*P*_):

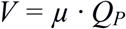

The growth rate is experimentally set by the chemostat dilution rate, here 0.3 d^−1^. The cellular P-quota (measured at 14.2 amol P cell^−1^) yielded a theoretical uptake rate of 4.3 ± 0.5 amol P cell^−1^ d^−1^. In comparison, the highest measured uptake rate was 1.3 ± 0.2 amol P cell^−1^ d^−1^, or about 30% of the theoretical value. This was the highest uptake rate measured for the culture and, consistent with results from the batch culture, indicated that HIMB114 growing under steady-state conditions with PO_4_^3-^ concentrations at 100 nmol L^−1^ was likely not using PO_4_^3-^ as a sole or primary P source for growth, and was instead meeting a large fraction of its P requirements from assimilation of organic P.

### Bacterial production

Bacterial production measurements were conducted on four consecutive days spanning the end of log phase into the early transition to stationary phase from the five batch culture experiments (Fig. 3A, Fig. S4). The mean per cell rate of leucine incorporation across all cultures grown on K-Bay standard medium was 0.042 ± 0.02 amol Leu cell^−1^ h^−1^ (mean ± s.d.; n=20), resulting in average cell-specific rates of production of 0.13 ± 0.07 fmol C cell^−1^ d^−1^ (mean ± s.d.; n=20) (Table 1).

**Table 1.**
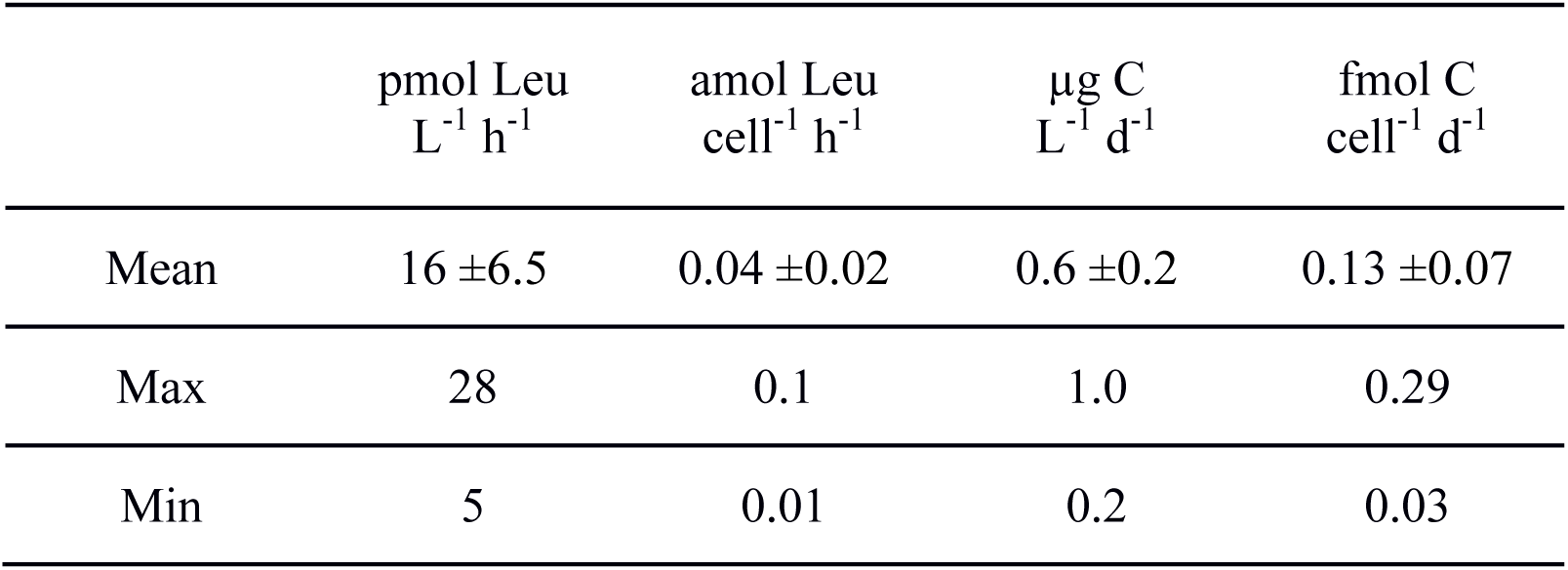
Mean (± standard deviation), minimum, and maximum bacterial production rates for SAR11 strain HIMB114 grown on sterilized K-bay seawater medium. Production (n=20) was measured in both batch and chemostat cultures by ^3^H-Leucine incorporation and converted to C units with a leucine to C conversion factor of 1.5 kg C mol Leu^−1^.

**Figure 3.**
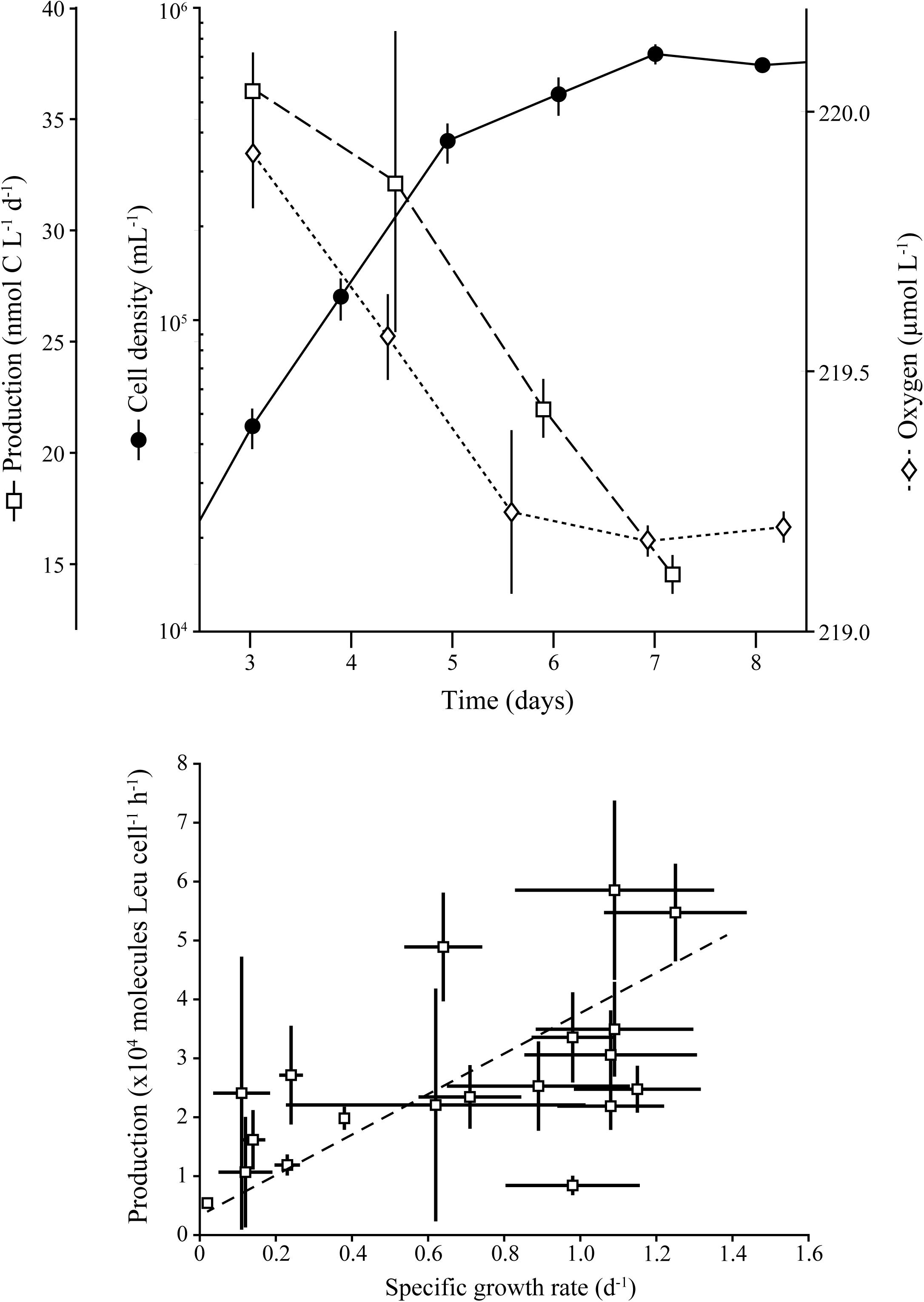
Production and respiration of strain HIMB114 during growth in batch culture. (A) Cellular growth (filled circles), bacterial production (^3^H-Leu; open squares), and dissolved oxygen concentrations (open diamonds) for a 10 L batch culture of HIMB114 measured throughout late exponential and into stationary phase (Table 2, experiment 1). (B) Bacterial production, expressed as molecules leucine per cell per hour over two-hour incubations, vs. daily specific growth rates for the same batch cultures of strain HIMB114, calculated by changes in cell densities between cultures sampled one day apart. The slope of the fit line between production and daily specific growth rate is 0.8 ± 0.3 (95% C.I.) ×10^6^ molecules Leu cell^−1^.

Bacterial production measurements for HIMB114 were relatively uniform, with a coefficient of variation of 40% across the 10 different cultures and 20 independent measurements grown on natural K-Bay seawater medium, despite being measured across a range of growth rates throughout exponential and early stationary phases of batch growth (linear correlation coefficient of 0.58, p-value 0.009) (Fig. 3B). However, in individual batch culture experiments, a decline in rate of cell division associated with entry into stationary phase was associated with a concomitant decline in production measured by leucine incorporation (Fig. 3A, Fig. S4), and the slope of the fit line for the plot of bacterial production vs. specific growth rate was positive (0.8 ± 0.3 (95% C.I.) ×10^6^ molecules Leu cell^−1^).

**Table 2.** Summary of experiments to measure respiration by the time dependent consumption of dissolved O_2_ in late-log phase batch cultures SAR11 strain HIMB114. Growth rates were calculated by linear least squares regression of cell density over the 2-day incubation periods. Note that Experiment 1 was conducted toward the transition out of log-phase and into stationary phase growth, with an associated flattening growth curve. Respiration was measured by linear least squares regression of Oxygen concentration with time over the 2-day incubations. Bacterial production values are 2-day means of daily, 2-hour Leucine incubations. Bacterial growth efficiency (BGE) was calculated as described in the text, assuming a respiratory quotient of 1 mol C: mol O_2_.

| Exp. | Growth rate | Abundance | Respiration |  | Production |  | BGE |
| --- | --- | --- | --- | --- | --- | --- | --- |
|  | Slope<br>±s.e.<br>(d<sup>-1</sup>) | Mean<br>[min, max]<br>x10<sup>8</sup> cells L<sup>-1</sup> | Mean<br>[min, max]<br>μmol O<sub>2</sub> L<sup>-1</sup> d<sup>-1</sup> | Mean<br>(±95% C.I.)<br>fmol O<sub>2</sub> cell<sup>-1</sup> d<sup>-1</sup> | Mean<br>[min, max]<br>μg C L<sup>-1</sup> d<sup>-1</sup> | Mean<br>(±95% C.I.)<br>fmol C cell<sup>-1</sup> d<sup>-1</sup> | Mean<br>[min, max]<br>(%) |
| 1 | 0.15 ±0.2 | 4.0<br>[3.5, 5.3] | 0.36<br>[0.10, 0.61] | 0.90 ±0.65 | 0.41<br>[0.30, 0.52] | 0.09 ±0.03 | 9<br>[5, 26] |
| 2 | 0.9 ±0.2 | 3.3<br>[1.9, 4.8] | 0.44<br>[0.27, 0.60] | 1.32 ±0.76 | 0.82<br>[0.70, 0.93] | 0.27 ±0.06 | 13<br>[10, 21] |
| 3 | 0.5 ±0.1 | 4.2<br>[3.3, 4.8] | 0.32<br>[0.14, 0.50] | 0.75 ±0.44 | 0.77<br>[0.66, 0.87] | 0.17 ±0.04 | 17<br>[11, 32] |

### Respiration

Rates of respiration were determined from three incubation experiments subsampled from the batch cultures (Table 2). Respiration was derived from linear regression fits to time course experiments in which the concentration of O_2_ was measured over two-day incubation periods (Figs. 3A and 4). Although extensive measures were taken to thoroughly acid clean and rinse the glass bottles, HIMB114 cells were only able to grow in the glass incubation bottles for a period of about two days and the consumption of O_2_ was only linear over this initial ∼two-day period (Fig. 4). Respiration rates showed high reproducibility over the three incubation experiments [0.37 ± 0.06 µmol O_2_ L^−1^ d^−1^ (mean ± s.d., n=3)] (Table 2) for cultures beginning with cell densities near 2 × 10^5^ mL^−1^ at the start of the incubations, and increasing on average 2.5 times over the two-day incubation period to about 5 × 10^5^ mL^−1^. Cell-normalized rates of respiration averaged ∼1 fmol O_2_ cell^−1^ d^−1^ when cultures were transitioning from exponential growth to stationary phase at a specific growth rate of approximately 0.5 d^−1^.

**Figure 4.**
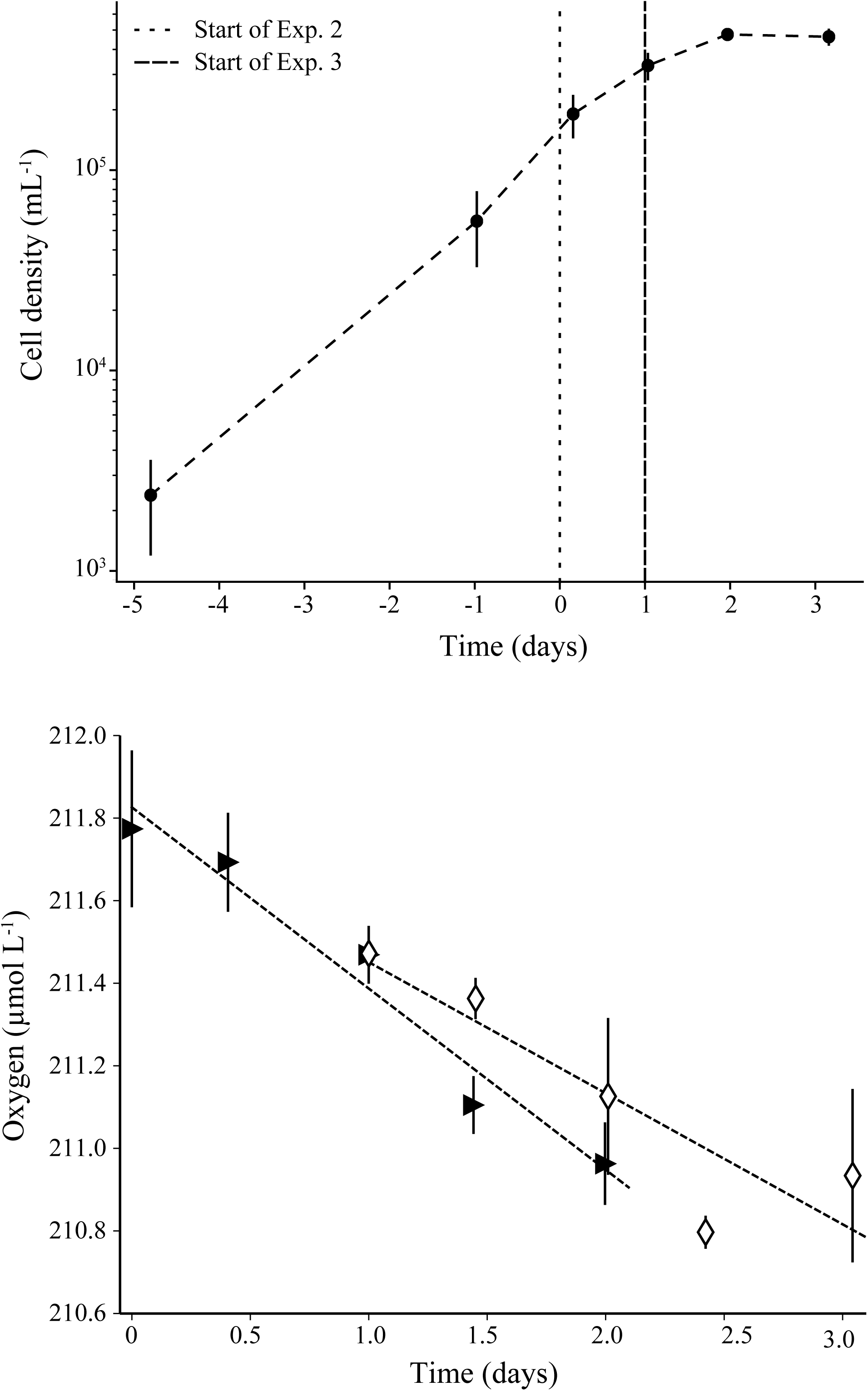
Respiration of strain HIMB114 during growth in batch culture. (A) Two oxygen respiration incubation experiments (experiments 2 and 3) started from a 10 L batch culture of strain HIMB114 during late exponential phase of growth. (B) Triangles and diamonds represent the mean oxygen concentration for each time point in the second and third incubation experiment, respectively (Table 2, experiments 2 & 3). Error bars represent the standard deviation.

**Table 3.**
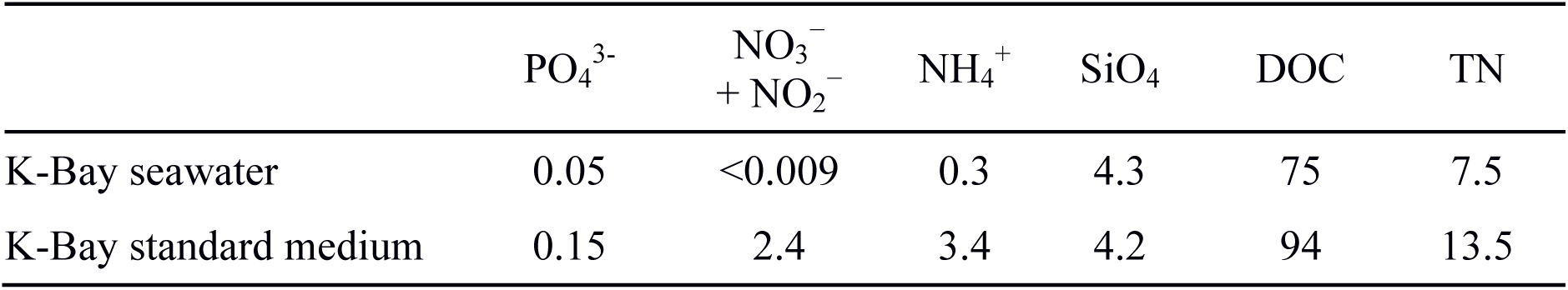
Nutrient concentrations (µmol L^−1^) for dissolved phosphate (PO_4_^3-^), nitrate plus nitrite (NO_3_^−^ + NO_2_^−^), ammonium (NH_4_^+^), silicate (SiO_4_), total organic C (TOC), and total nitrogen (TN) for natural Kaneohe Bay (K-Bay) seawater with no additions, and the standard SW based medium used to grow HIMB114 in both batch and chemostat cultures.

Rates of respiration were also derived based on changes in total organic C (TOC) within HIMB114 cultures over several days, measured at the start and end of the incubations of six replicate 10 L batch cultures of HIMB114 (Fig. S1). The initial TOC concentration in the media was 83 ± 2 µmol C L^−1^ (mean ± s.d., n=6), while the final TOC concentration sampled 9 days later was 79 ± 3 µmol C L^−1^ (mean ± s.d., n=6), resulting in a mean drawdown of TOC over the 9-day incubation of 4 ± 4 µmol C L^−1^ (mean ± s.d., n=6). This is equivalent to approximately 5% of initial TOC. The resulting average rate of respiration was 0.44 ± 0.44 µmol C L^−1^ d^−1^. This rate is very similar to the mean rate of O_2_ consumption (based on the two day incubation period), assuming a respiratory quotient of 1 mol C: mol O_2_, of 0.37 ± 0.06 µmol C L^−1^ d^−1^ (mean ± s.d., n=3).

### Bacterial growth efficiency

By combining rates of bacterial production with the measured rates of respiration we were able to estimate bacterial growth efficiency (BGE) for the HIMB114 batch cultures. BGE is defined as the ratio of the C production rate to the total C demand, which is the sum of production and respiration:

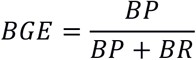

Combining these rate measurements resulted in a mean BGE of 13% with a 95% confidence interval of 10 – 21% estimated by a Monte Carlo simulation study (Table 2).

## Discussion

Steady state chemostat growth provides an ideal system to investigate the physiology and cellular properties of model microorganisms. The chemostat system allows the investigation of cellular physiology under controlled growth rate conditions, something unachievable using batch cultures. In this case, the steady state growth achieved through the chemostat allowed us to simultaneously calculate the theoretical P demand as well as determine the actual PO_4_^3-^ uptake rate at a set rate of growth. Despite the inability to grow strain HIMB114 under P-limiting conditions, these experiments suggest that this isolate relies heavily on sources of P other than PO_4_^3-^ when grown on a natural seawater minimal medium. This finding is particularly intriguing considering that HIMB114 has a complete high-affinity PO_4_^3-^ transport system (19), and so should have the full capacity to transport PO_4_^3-^ under dilute conditions. While inorganic PO_4_^3-^ is generally considered the preferred P source for marine bacteria (14), oligotrophic marine environments such as Kaneohe Bay in the tropical Pacific Ocean where strain HIMB114 was isolated typically have DOP concentrations an order of magnitude above inorganic phosphate concentrations (20). Thus, the ability to utilize components of the DOP pool to attain P may be competitively advantageous. In both batch and chemostat culture conditions, HIMB114 appeared to utilize an undetermined component of the DOP pool to meet its P growth demands, even when PO_4_^3-^ was amended to the media. Which component(s) of the DOP pool were utilized remains to be determined. One potential class of DOP compounds receiving recent attention are phosphonates, which are organic phosphonic acid derivatives containing a C-P bond and which make up 25% of the high molecular weight DOM pool (21). Evidence of the widespread distribution of genes for the transport and metabolism of phosphonates has been reported in marine microorganisms (22-24) including SAR11 (Table S1) (17, 19), and there is precedent for the PO_4_^3-^-independent utilization of phosphonates in marine systems (25). In laboratory experiments with a defined growth medium, SAR11 subgroup Ia strain HTCC7211 was shown to utilize phosphonates as a source of P for growth (12). While the genome of strain HIMB114 encodes for a complete phosphonate transport system similar to that of HTCC7211, it encodes a unique and sparse complement of genes for phosphonate metabolism (Table S1).

At 14.2 amol P cell^−1^, the measured P cell quota for HIMB114 is close to that determined for SAR11 subgroup Ia strain HTCC7211 (10.9 amol P cell^−1^) grown on a P-limiting, defined medium (12). Using transmission electron microscopy coupled with X-ray microanalysis, Gundersen and colleagues found an empirical biovolume (*V*) power law relationship for cell P quotas of 126*V*^0.937^ amol P cell^−1^, measured for 84 bacterial cells with mean cell volume 0.08 µm^3^ (range 0.001 to 2.0 µm^3^) (26). Using this power law for HIMB114 cells suggests a cellular quota of 13 amol P cell^−1^, similar to the value measured in our study. At 1.237 Mbp and 2 P-atoms per base pair, the small genome of HIMB114 yields a P content of 4.1 amol P cell^−1^. This calculation suggests DNA alone accounts for approximately 29% of the cellular P-quota, a finding roughly 2 to 3 times the 10-15% cellular P traditionally considered accounted for by DNA in a bacterial cell (27). Another significant pool for P is likely phospholipids, which have previously been measured to contain 2.5 amol P cell^−1^ for strain HIMB114 grown in PO_4_^3-^ replete seawater media (28). Thus, nearly half of the P-quota of the cell can be accounted for by only two macro-molecular components: DNA and phospholipids. While not quantified in HIMB114 or other SAR11 cultures, RNA is typically the dominant contributing molecular pool to cellular P; typical total RNA to DNA mass ratios are infrequently below 2:1 (mass RNA: mass DNA), even for slowly growing bacteria (29). At a lower limit of 1:1 RNA:DNA, an additional 4.1 amol P cell^−1^ can be accounted for by inclusion of cellular RNA pools. Hence, HIMB114 appears to have a similar cellular P concentration (0.16 M P) compared to other marine bacteria (0.1 to 0.2 M P) (30), and it would be difficult to further reduce this P demand unless these cells were able to substitute P-free lipids for phospholipids as has been demonstrated for SAR11 subgroup Ia strain HTCC7211 (31). However, no genetic capacity for phospholipid substitution analogous to that found in the genome of strain HTCC7211 is apparent in the HIMB114 genome.

At 0.05 µm^3^, the peak of the distribution for cell volume of strain HIMB114 was consistent with the recently reported range of 0.015 to 0.058 µm^3^ for SAR11 subclade Ia isolates HTCC1062 and HTCC7211 (3). However, the mean value of the cell volume distribution (0.09 µm^3^) for HIMB114 was larger than anticipated, which can be at least partially attributed to elongated and chains of cells that increase in frequency as HIMB114 enters into stationary phase (Fig. 1). This phenomenon has been observed previously for SAR11 subgroup Ia strain HTCC1062 (5) and thus may be a broadly distributed, growth stage-dependent feature of SAR11 that has the potential to confound models and other measurements that rely on an average cell size or that are normalized per cell. For example, this phenomenon may contribute to the variability observed between bacterial production measured by leucine incorporation and cellular growth rate during the transition to stationary phase (Fig. 3).

The genome and the membrane envelope are two essential components of a cell that cannot be continuously scaled down with cell size (6), and hence represent increasing fractions of total cell volume, or mass, for cell volumes below 0.05 µm^3^ (Fig. S5). This constrains the lower limit for a bacterial cell volume to about 0.004 µm^3^. We therefore propose a potential trade-off between nutrient acquisition strategies and P growth requirements for small cells. In oligotrophic, nutrient-limited environments, a high surface area to volume ratio should increase a cells ability to compete for dilute nutrients, giving small cells a distinct competitive advantage. Yet, there is an opposing force balancing this trend toward smaller cell size, namely an increasing P requirement relative to cell mass necessary to maintain a given growth rate. This trade-off is reflected in the importance of P-sparing strategies employed by oligotrophic picocyanobacteria such as *Prochlorococcus* (32), and the noted prevalence and diversity of P acquisition and metabolism related genes found to be important across large ocean ecosystem regimes (15, 17). In addition to membrane lipid renovation (31), strategies employed by very small, diverse, and successful bacteria of the SAR11 clade to sustain cellular P demands and otherwise maintain sufficient net growth rates to numerically dominate surface marine waters will no doubt continue to provide interesting discoveries.

The exceedingly high C to N stoichiometry of near 40±14:1 (mol C: mol N) is well outside normally reported ranges for bulk marine particulate organic matter or C:N ratios of flow cytometrically sorted natural planktonic populations (max 24.4, mean 9.4±3.6, n=277) (33). Moreover, the resulting C: N: P cellular stoichiometry for HIMB114 would approach 300:7:1 (mol C : mol N : mol P), a finding inconsistent with previous estimates for members of the SAR11 clade. Such results are primarily driven by the exceedingly large C cell quota measured for HIMB114 in this study (50 ± 47 fg). The N cell quota for HIMB114 (1.4 ± 0.9 fg) is slightly lower than the minimum N cell quota of 1.6 fg N for natural bacteria reported by Fagerbakke, Heldal & Norland, (30). Using the biovolume power law regression from Gundersen et al., (26) to derive cellular N quotas results in 2.4 fg N for a cell volume of 0.09 µm^3^ (mean volume measured in the current study); the same relationship yields a cellular C quota of ∼13 fg C. Tripp and colleagues previously estimated that SAR11 subgroup Ia strain HTCC1062 contained 5.8 fg C cell^−1^ for cells of biovolume 0.035 µm^−3^ (7), which scales to 14.9 fg C cell^−1^ for a cellular volume of 0.09 µm^3^. Similarly, Cermak and colleagues estimated that cellular C quotas varied between 12 to 16 fg for SAR11 strains HTCC1062 and HTCC7211 (34), which would scale to 24 to 48 fg C for HIMB114 when accounting for differences in cell volume. Such comparisons suggest the measurements of cellular C from the current study are overestimates. Although it remains unclear what factors may have contributed to these results, such findings may reflect the poor filtration retention efficiency of the filters utilized for these measurements. Regardless, accurate quantification of cellular C content of SAR11 cells remains imperative for future research efforts.

Although there are no published bacterial production measurements for any axenic SAR11 cultures, we can compare our values to measurements from planktonic marine ecosystems where SAR11 often dominate. The observed mean leucine incorporation rate from this study (4.2 ×10^−8^ pmol Leu cell^−1^ h^−1^) is very close to that of the natural community mean dark leucine incorporation rates, normalized to non-pigmented cell counts, for station ALOHA in the North Pacific subtropical gyre of 5×10^−8^ pmol Leu cell^−1^ h^−1^ (35), and falls at the lower end of the range measured for natural surface seawater communities along a transect off the Oregon coast (0.39–4.7 ×10^−7^ pmol Leu cell^−1^ h^−1^) (36). Malmstrom and colleagues measured the contribution of naturally occurring SAR11 populations to bulk ^3^H-leucine incorporation rates using a combination of microautoradiography and fluorescence *in situ* hybridization (Micro-FISH) in the Northwest Atlantic Ocean (37). These authors reported that SAR11 accounted for a large fraction (50%) of the bulk leucine incorporation rates in surface waters, where they represented 25–35% (2–4×10^8^ cells L^−1^) of the picoplankton population. The resulting SAR11-specific C production rates were estimated to be from 0.5 µg C L^−1^ d^−1^ for an open-ocean Gulf Stream site, increasing to about 3 µg C L^−1^ d^−1^ for a coastal location. del Giorgio and Cole compiled published marine ecosystem bacterial production measurements and reported global mean bacterial production rates of 2.41 ± 0.33 µg C L^−1^ d^−1^ for the coastal ocean to 0.37 ± 0.054 µg C L^−1^ d^−1^ for the open ocean (38). In the current study, the bulk C production rate measured for HIMB114 cultures was 0.6 ± 0.2 µg C L^−1^ d^−1^, similar to those estimated by Malmstrom and colleagues (37) and typical of open-ocean, oligotrophic values reported by del Giorgio and Cole (38).

Leucine incorporation rates are used as a standard proxy for biomass production under the assumption that protein is a major constituent of cell biomass, and that leucine represents a relatively stable proportion of bacterial protein (39). This allows consistent comparisons of protein synthesis rates, and thus biomass production, across the wide spectrum of bacterial species capable of taking up leucine. It was somewhat surprising then to find that bacterial production measured by leucine incorporation varied little across a range of growth rates for HIMB114, with only a weak correlation between the two measures. While this could be due to the low precision (typically 15%) for both cell counts and leucine incorporation measures, it is also possible that protein production rates and cell division rates are uncoupled at short time scales under non-linear batch growth.

We also measured O_2_-based rates of respiration, with rates averaging ∼0.4 µmol O_2_ L^−1^ d^−1^. These low rates of respiration were highly reproducible, with a coefficient of variation of 16% between replicate incubations. Moreover, the measured O_2_-based respiration measurements agreed with the measured drawdown of organic carbon over the course of the incubations, which together with measured rates of bacterial production yielded estimates of BGE near 13%. Such results suggest HIMB114 grows at similar efficiencies as other marine heterotrophic bacteria, despite features such as an exceptionally small, streamlined genome that might be expected to enable more efficient growth.

Steindler and colleagues have published the only other dissolved oxygen measurements from a SAR11 culture, wherein SAR11 subgroup Ia strain HTCC1062 cultures were measured using an oxygen optode (4). Although rates of respiration were not explicitly calculated in that study, over the initial 69 h period of incubation concentrations of O_2_ declined by approximately 100 µmol O_2_ L^−1^, equivalent to a rate of O_2_ consumption of ∼35 µmol O_2_ L^−1^ d^−1^. Cell densities in the experiments of Steindler and colleagues were three orders of magnitude higher than the densities of our experiment; when normalized to cell density, the rate of respiration for HTCC1062 was approximately 0.35 fmol O_2_ cell^−1^ d^−1^, approximately one third of the rate observed for strain HIMB114.

## Conclusions

Though we were unable to create a state of P-limited growth or to determine what may be limiting HIMB114 when grown on a minimal seawater medium, we were able to rule out many of the common C, sulfur, and specific amino acid growth substrates that have been shown to enhance growth for other SAR11 cultures and permit their growth in a defined, artificial seawater growth medium (5, 7, 8, 11, 40). While this implies caution in extrapolating the results of culture-based studies from specific SAR11 isolates to the SAR11 lineage as a whole, it also suggests that exciting metabolic features that distinguish populations, ecotypes, and major SAR11 sub-lineages await characterization. One such feature, uncovered by using a continuous culture of HIMB114, is the apparent inability of this strain to fulfill its cellular P-demand through the uptake of PO_4_^3-^ alone. Our findings support the idea that at least some members of the SAR11 clade rely on organic P to support growth, which is also supplemented by the use of cellular P-sparing adaptations such as lipid renovation (1). Despite potential methodological issues with the measurement of cellular C content, the N and P cell quotas, production, respiration, and cell size measurements reported here provide new information for scientists and modelers interested in understanding the impact of SAR11 cells on the ecology of the global ocean.

## Materials & Methods

SAR11 strain HIMB114 was previously isolated from Kaneohe Bay on the northeastern shore of the island of Oahu in the tropical Pacific Ocean using a dilution-to-extinction approach (2, 41). It is a member of subclade IIIa that, based on genome comparisons, exhibits genus-level divergence from the comparatively well-studied members of subgroup Ia (i.e. ‘*Candidatus* Pelagibacter’) (19, 42, 43). HIMB114 would not grow in defined artificial seawater-based media previously published for SAR11 (5, 40), nor could we enhance its cellular yield by previous organic carbon, vitamin, and nutrient additions that have proven successful for other SAR11 strains (7, 8, 11) (data not shown). Thus, all experiments were performed in natural seawater-based minimal media with seawater collected from the southern basin of Kaneohe Bay (21° 26.181’ N, 157° 46.642’ W). To make growth media, 200 L of surface seawater was filtered through pre-rinsed (10 L sterile water followed by 10 L seawater) 0.1 µm pore-sized polyethersulfone (PES) membranes (AcroPak 1000; Pall Corp., Port Washington, NY, USA) into clean 10 L polycarbonate (PC) carboys. Individual 10 L batches of seawater were subsequently autoclaved for 2.5 hours (h) at 121 °C and allowed to cool. For both batch and chemostat media (media termed “K-Bay”), the seawater base was amended with nitrate (3 µmol L^−1^ NaNO_3_), NH_4_^+^ (3 µmol L^−1^ NH_4_Cl), PO_4_^3-^ (0.1 µmol L^−1^ KH_2_PO_4_), and a vitamin stock solution added at 10^−5^ dilution (10^−6^ dilution for the chemostat medium) (2). All chemicals were BioUltra grade (MilliporeSigma, St. Louis, MO, USA). The vitamin stock solution contained B1 (thiamine hyrdochloride, 1 g L^−1^); B3 (niacin, 0.1 g L^−1^), B5 (pantothenic acid, 0.2 g L^−1^); B6 (pyridoxine, 0.1 g L^−1^); B7 (biotin, 1 mg L^−1^); B9 (folic acid, 2 mg L^−1^); B12 (cyanocobalamin, 1 mg L^−1^); myo-inositol (1 g L^−1^); and PABA (4-aminobenzoic acid, 0.1 g L^−1^). Following nutrient additions, small amounts of autoclaved-sterile milli-Q deionized-water was added to the K-Bay media to replace water lost as a result of autoclaving, achieving a final salinity of 32. The media were then sparged with CO_2_, followed by air, through three in-line Whatman (GE Healthcare Life Sciences, Chicago, IL, USA) vent filters (HEPA 0.3 µm glass fiber to 0.2 µm PTFE to 0.1 µm PTFE) to restore the inorganic C chemistry and to bring the media pH to between 8.0 and 8.1, and stored at 4 °C until use. HIMB114 cultures were grown in batch at 26 °C under low light (33 µmol quanta m^−2^ s^−1^) and a 12/12 light/dark cycle in volumes ranging from 100 mL to 10 L, as well as a 4 L continuous culture chemostat system (described below).

### Chemostat

For continuous culture growth, a custom-built 4 L PC chemostat was constructed using a narrow mouth 4 L PC bottle, 4-port Teflon threaded cap, PC Luer connection fittings, and silicone tubing for the inflow of growth medium, culture overflow, air bubbling, and culture sampling ports. The chemostat was kept under positive pressure by bubbling with 0.1 µm-filtered air, which served to keep the culture well-mixed as well as provide positive pressure for culture sampling. Media was pumped from a 10 L PC carboy continuously at 0.85 mL min^−1^ for a target dilution rate of 0.3 d^−1^. Overflow was continuously removed into a PC bottle used as an overflow container. To start the continuously growing culture, the chemostat was filled to 2 L with the K-Bay chemostat medium (Table 2), inoculated with 5 mL of a growing HIMB114 culture, and allowed to grow in batch, where it reached an exponential growth rate of 0.75 d^−1^ for 10 days before media in-flow was started (Fig. 5). After reaching the full 4 L chemostat volume, cell densities stabilized at 7×10^5^ mL^−1^ after approximately 5 days, and remained in continuous culture for 12 days or about 5 doubling times with continuous media inflow and culture overflow (Fig. 5). The culture was grown in the chemostat for a total of 40 days; however, following the connection of the third 10 L batch of new K-Bay chemostat medium at day 30, cell densities slowly declined to 3.5×10^5^ mL^−1^ by the end of the 40 days when the chemostat was turned off (Fig. 5).

**Figure 5.**
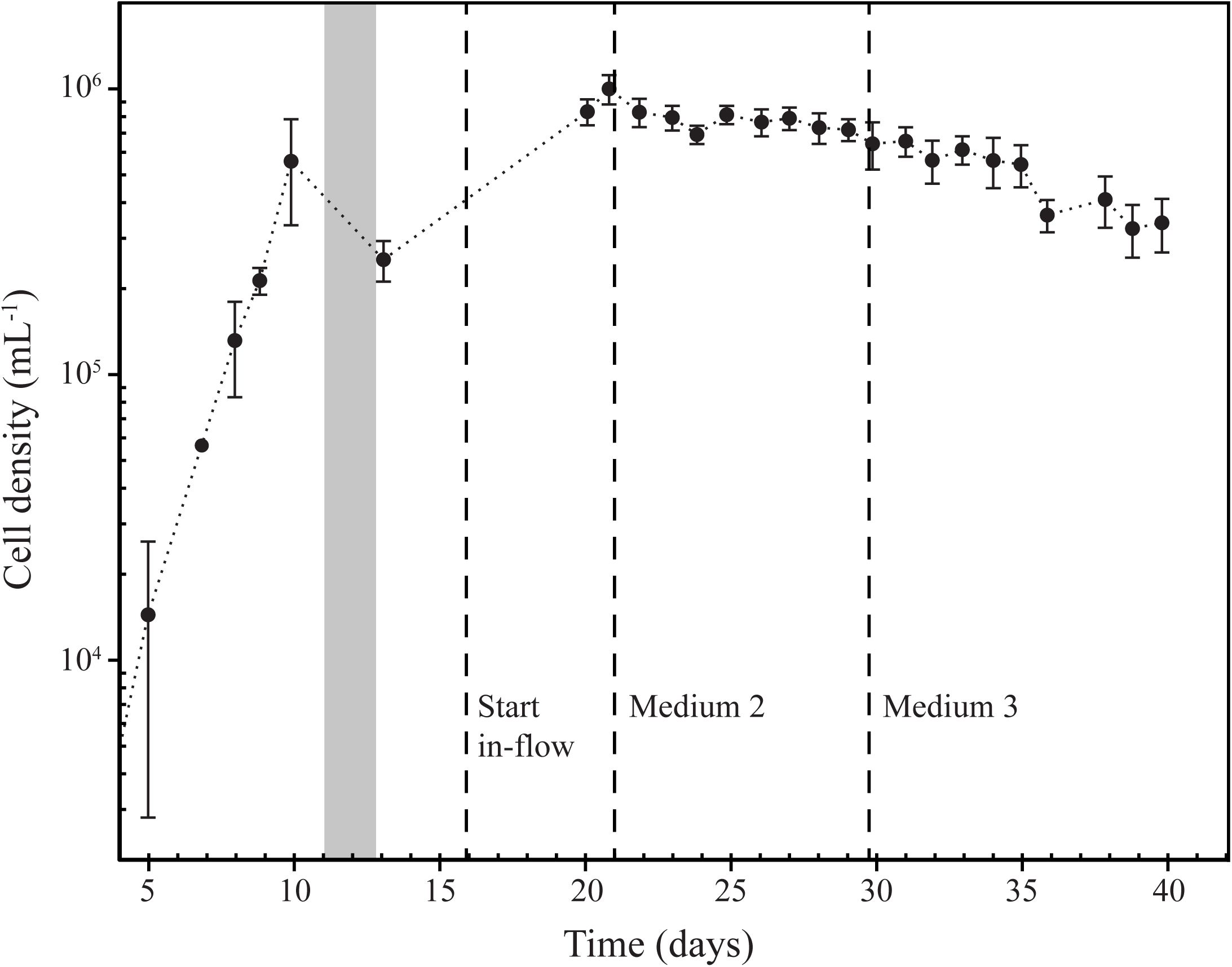
Chemostat continuous culture of SAR11 strain HIMB114. HIMB114 cells growing at 0.3 d^−1^ in chemostat continuous culture in natural seawater media at 26 °C. Standard deviation of cell counts are indicated by error bars. The grey box indicates the time when filling the chemostat, while dotted lines indicate when media in-flow started and new 10 L media reservoirs were connected.

### Cell enumeration and image analysis

Culture cell counts were made by 4’,6-diamidino-2-phenylindole (DAPI) staining and subsequent epifluorescence microscopy. Depending on culture density, between 2 to 10 mL culture samples were fixed with 20% electron microscopy grade paraformaldehyde solution (Electron Microscopy Sciences, Hatfield, PA, USA) to a final fixative concentration of 0.4%, and stored at 4 °C overnight. DAPI was subsequently added to a final concentration of 5 µg L^−1^ and incubated in the dark at room temperature for at least 20 minutes. Stained samples were filtered onto 25 mm diameter, 0.2 µm pore-sized, black Nuclepore (GE Healthcare Life Sciences) or Isopore (MilliporeSigma) PC membranes, with a 0.8 µm pore-sized GN-4 (Pall Corp.) mixed cellulose ester backing filter. Filters were allowed to air dry for 15 minutes and either stored frozen (−20 °C) or mounted in high-viscosity immersion oil on a glass slide for microscopic enumeration. At the volumes filtered, the precision of epifluorescence microscope cell counts were between 10% to 20% for densities above 10^4^ mL^−1^. Cell number and size information (including cell lengths and widths) were calculated by image detection software from DAPI stained epifluorescence images captured with a Retiga EXi FAST1394 camera (QImaging, Surrey, BC, Canada) at 1000x magnification.

Cell morphology was also visualized via scanning electron microscopy. HIMB114 cells grown in K-bay media to early stationary phase were fixed with gluteraldehyde (20%; Electron Microscopy Sciences), filtered onto 0.2 μm pore-sized PC membranes (Nuclepore), washed with sodium cacodyolate buffer, post-fixed in osmium tetraoxide, and subjected to sequential ethanol dehydration, critical point drying with CO_2_, and coating with gold/palladium. The preps were viewed on a Hitachi S-4800 Field Emission Scanning Electron Microscope with Oxford INCA X-Act EDS System.

### PO_4_^3-^ uptake

Rates of PO_4_^3-^ uptake were measured in both batch and continuous culture using ^33^P-radiotracer methods (18). In brief, between 10–50 mL aliquots of growing culture were sampled into 50 mL PC tubes and spiked with ^33^P-orthophosphoric acid (158 Ci mg^−1^; PerkinElmer, Waltham, MA, USA) to a specific activity of 50 µCi L^−1^. ^33^P-labeled cultures were incubated (typically for 4 to 8 h) under identical growth conditions to the parent cultures. Sample time points were collected by low-vacuum filtration of 5 mL onto 0.2 µm pore-sized, 25 mm diameter PC Nuclepore membranes, each filter having been pre-saturated with unlabeled PO_4_^3-^ by the addition of 1 mL of high PO_4_^3-^ (0.1 mmol L^−1^ PO_4_) seawater to each filter prior to sampling (Suppl. Info.). Following filtration of ^33^P-labeled samples, filters were rinsed with 10 mL of 0.2 µm-filtered seawater. Along with the culture samples, 0.2 µm-filtered seawater controls amended with ^33^P-radiotracer served as non-biological blanks; these blanks were incubated and processed identical to samples. Phosphate uptake kinetics were also conducted for nine treatments of increasing PO_4_^3-^ concentration by the addition of 1 – 20 µL of concentrated unlabeled phosphate stocks (0.1 – 1 mmol L^−1^ KH_2_PO_4_) to 11 mL chemostat culture samples. The treatments and filtered seawater controls were subsampled (2 mL) at 5 time points over 22 hours of batch growth.

### Bacterial production

Bacterial production was estimated based on the incorporation of tritiated-leucine (^3^H-Leu) into protein using small volume (1.5 mL) sample incubations based on the microcentrifugation method (44) (Suppl. Info.). Leucine incorporation rates were converted into C production rates using a standard 1.5 kg C mol Leu^−1^ conversion factor (36).

### Oxygen respiration

Respiration rates were measured in batch cultures of strain HIMB114 based on time-dependent changes in oxygen to argon (O_2_/Ar) ratios measured by membrane inlet mass spectrometry (MIMS) (45) (Suppl. Info.). Cultures growing in late exponential phase in 10 L PC carboys were siphoned using silicone tubing into 70 mL clear glass serum bottles, allowed to overflow, capped with Teflon lined rubber stoppers, and crimped sealed. The glass bottles were extensively cleaned with milli-Q DI-water and 10% hydrochloric acid, and finally autoclaved while filled with milli-Q DI-water before use. Sample bottles were filled in triplicate for each time point, with 5 time points sampled over a two-day period, and in one case a four-day period. Bottles were incubated under the same temperature and light conditions as the original cultures, and either run immediately at each time point or killed by syringe addition of 100 µL of saturated mercuric chloride solution and analyzed at the end of the incubations.

### Monte Carlo simulation

A Monte Carlo simulation study was conducted to quantify statistical errors in the oxygen-based respiration, bacterial production, and BGE measurements. Simulated data for O_2_ concentrations and leucine incorporation rates from the respiration and production experiments were generated by sampling (N=10,000) from independent, normal distributions using sample means and variances based on experimental replicate measurements. Linear regression slopes were computed on the simulated O_2_ concentration samples with time to obtain simulated O_2_ respiration rates separately for each of three incubation experiments, with regression slopes bounded by zero (i.e., simulated data was prevented from indicating net O_2_ production with time). To convert from O_2_ and leucine units into carbon units, no uncertainty was assumed in the conversion factors, as we were attempting to estimate the statistical error from our measurement replication. BGE was calculated as indicated above, and 95% confidence intervals (2.5% and 97.5% quantiles) were calculated for each measurement by experiment (Table 2) and for the final reported mean BGE measure.

### Dissolved nutrients

Samples for dissolved inorganic nutrients and TOC analyses were taken from both the original medium as well as the final spent medium at the end of culture incubations. Samples were collected in acid washed, DI-water rinsed plastic (inorganic nutrients) or glass (TOC) containers, and stored frozen until analysis. Dissolved inorganic nutrient samples were analyzed on a Analytical Segmented Flow Injection AutoAnalyzer AA3 HR (SEAL Analytical Inc., Mequon, WI) for the determination of PO_4_^3-^, NH_4_^+^, nitrate + nitrite (NO_3_^−^ + NO_2_^−^), silicate (SiO_4_), and total N. Samples for TOC were acidified and O_2_ purged to remove inorganic C, and measured using high temperature catalytic oxidation on a Shimadzu TOC-L (Schimadzu Scientific Instruments Inc., Columbia, MD).

### Cellular elemental analysis

Six individual 10 L cultures of HIMB114 were grown in batch for the purpose of collecting 20 L of cultured cells onto triplicate Advantec GF-75, 25 mm diameter glass fiber filters (Sterlitech, Kent, WA, USA), with a nominal pore size of 0.3 µm, for subsequent measurements of cellular C and N quotas. Filters were dried, pelleted, and analyzed using an elemental analyzer (CE440 elemental analyzer, Exeter Analytical, North Chelmsford, MA, USA). The batch cultures were filtered by slowly pumping the cultures from 10 L carboys into large volume filter towers containing combusted 25 mm GF-75 filters; the filtrate was retained in separate 10 L collection carboys for subsequent microscopic analyses to assess the cellular retention efficiency of the filters. A total volume of 30 L was filtered through each membrane filter: 20 L from two separate 10 L cultures, and 10 L of culture filtrate containing cells that passed through the first filtration. The cell retention efficiencies of the filters declined in each successive 10 L round, from a mean retention of 37% in the first round, down to 9% in the final third round of filtration from the filtrate. The overall cell retention rate for the full filtration procedure was 40%, resulting in an average of 5 ± 2 ×10^9^ cells (mean ± s.d.; n=3) on each filter. Preliminary tests indicated that procedural blanks were necessary to account for adsorption of non-cellular dissolved C and N onto the filter (Suppl. Info.).

For the determination of cellular P, 4 to 5 L of culture were collected by peristaltic pump filtration at a flow rate of 8 mL min^−1^ onto 0.2 µm pore-size, 47 mm diameter PC Nuclepore filters. All filtrations occurred in a walk-in cold room at 4 °C for 8 to 10 h. Procedural blanks of spent media were also made by filtering 50 mL of 0.2 µm media filtrate, that is, the same media in which the cultures were grown with cells removed, onto the 0.2 µm, 47 mm-diameter PC filters. Following filtration, filters were placed in acid-cleaned glass test tubes, covered with combusted aluminum foil and stored at −20 °C until analysis. Cellular P was quantified by a modification of the high temperature combustion, colorimetric molybdate method (46) (Suppl. Info.).

## Acknowledgements

We thank E. Omori and K. Manoi for laboratory assistance, the laboratory of Craig Carlson at the University of California, Santa Barbara for TOC measurements, and D. Karl for early review and feedback on this research. Nutrient samples were analyzed by the SOEST Analytical laboratory at the University of Hawaii at Manoa. This research was supported by funding from National Science Foundation grant OCE-1538628 to MSR and the Center for Microbial Oceanography: Research and Education (C-MORE, NSF Science and Technology Center award EF-0424599). SF was funded by a C-MORE fellowship. This is SOEST contribution xxxx and HIMB contribution xxxx.

## Supplemental Material

### Supplemental Methods

**Table S1.** Presence of genes for phosphorus uptake and metabolism in selected publicly available SAR11 genomes sequenced from cultivated strains.

| Strain | Lineage | Source | pstS<br>phosphate<br>transport | phoBU<br>phosphate<br>regulation | phnCDE<br>phosphonate<br>transport | phnAGHI<br>JKLMNX<br>phosphonate<br>metabolism | ppx/ppk<br>polyphosphate<br>metabolism |
| --- | --- | --- | --- | --- | --- | --- | --- |
| HIMB114 | IIIa | Subtropical<br>North Pacific | + | + | + | phnAX | - |
| HIMB58 | IIb | Subtropical<br>North Pacific | + | + | - | - | - |
| HIMB59 | V | Subtropical<br>North Pacific | + | + | + | phnAX | - |
| HIMB5 | Ia | Subtropical<br>North Pacific | + | + | - | - | - |
| HTCC7211 | Ia | Sargasso Sea,<br>Atlantic | + | + | + | + | + |
| HTCC9565 | Ia | Temperate<br>North Pacific | - | phoH | - | - | - |
| HTCC1002 | Ia | Temperate<br>Coastal Pacific | - | phoH | - | - | - |
| HTCC1062 | Ia | Temperate<br>Coastal Pacific | + | + | - | - | - |

**Figure S1.**
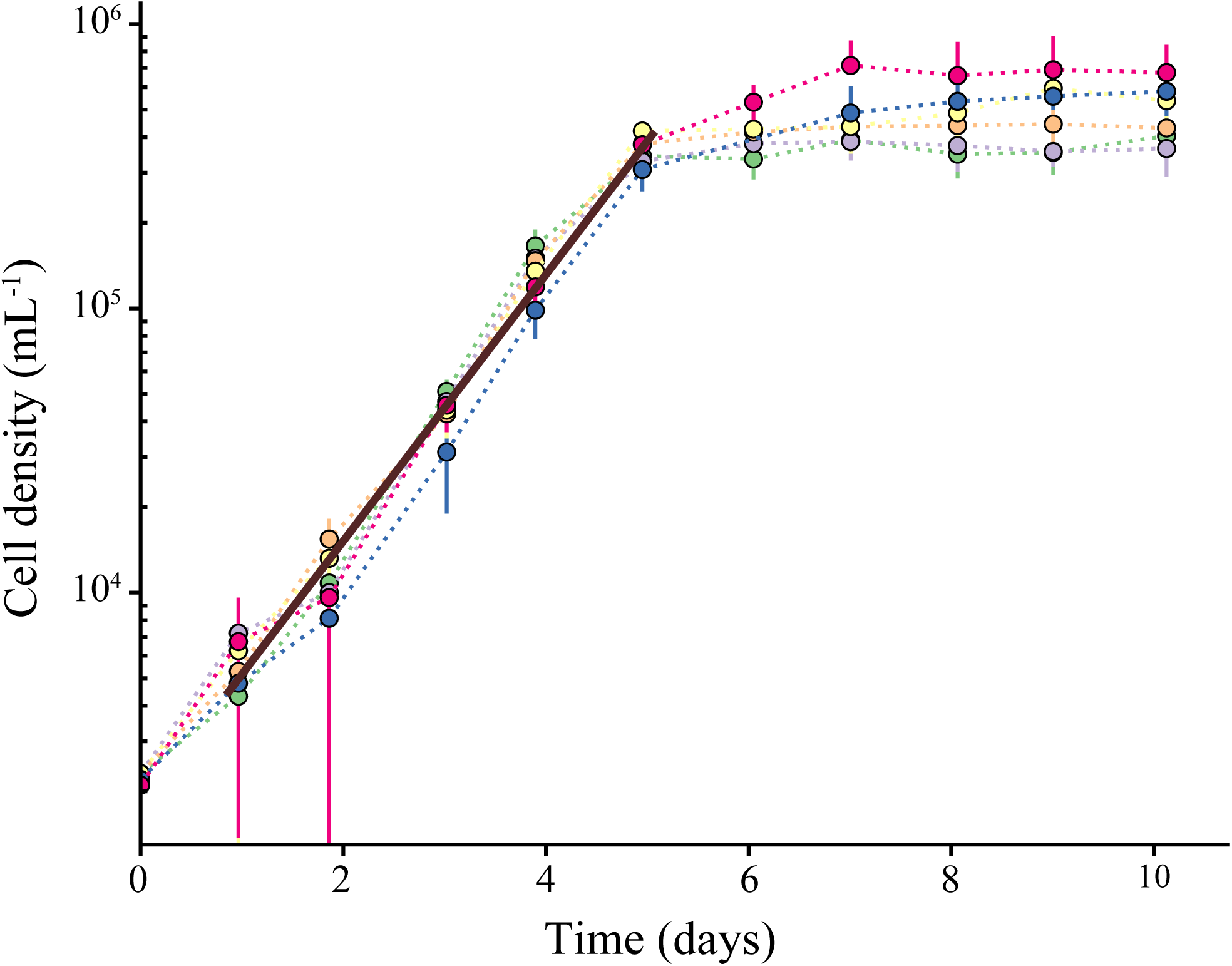
Batch culture growth curves for six replicate 10 L cultures of HIMB114. Lower error bars represent the standard deviation among count fields (typically 10-15), while upper error bars represent the cell density if morphologies representing multiple cells are converted to single cell units. The mean regression slope for exponential specific growth rate for the cultures is 1.08 (s.d. 0.03) d^−1^. Strains were grown in using standard media at 26 °C.

**Figure S2.**
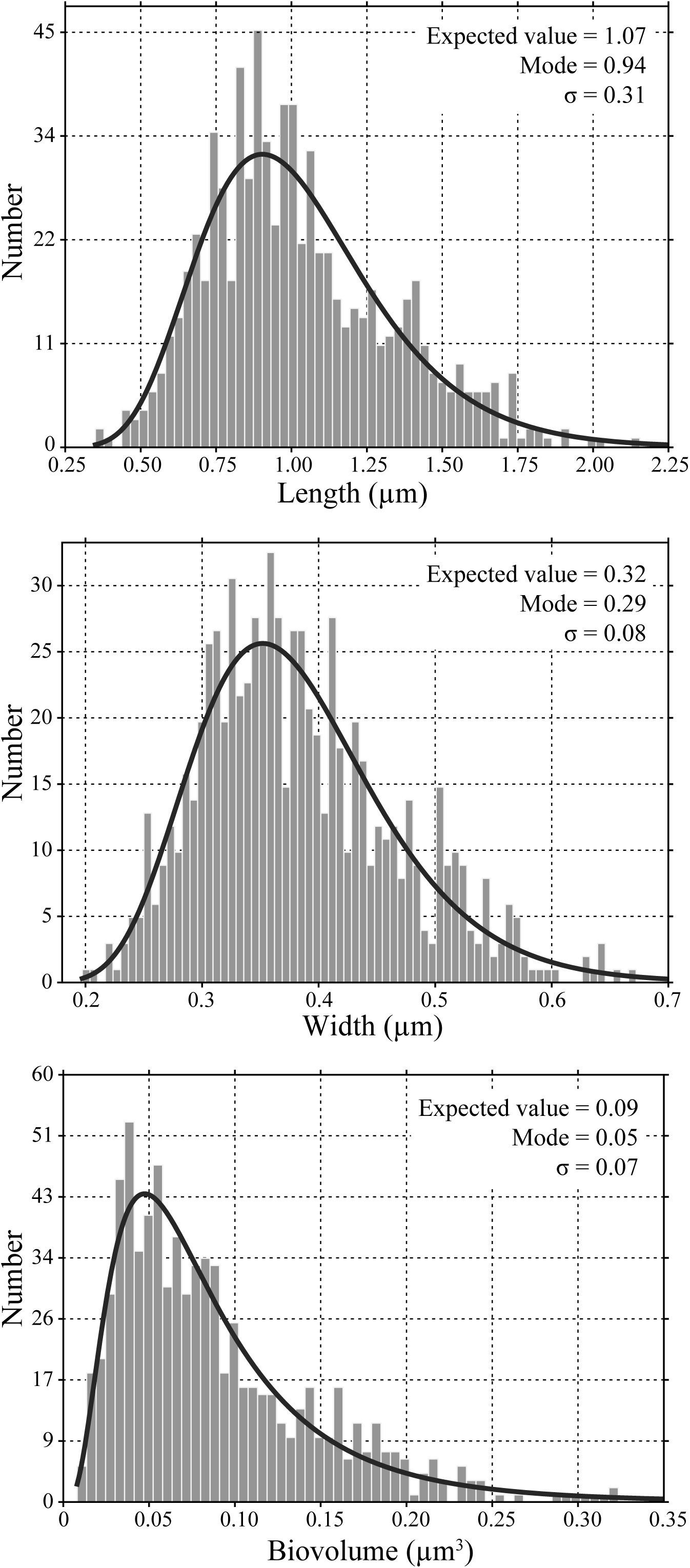
Cell size analysis of SAR11 strain HIMB114. Distribution of (A) length, (B) width, and (C) biovolume for N=769 cells, with fit to log-normal distribution.

**Figure S3.**
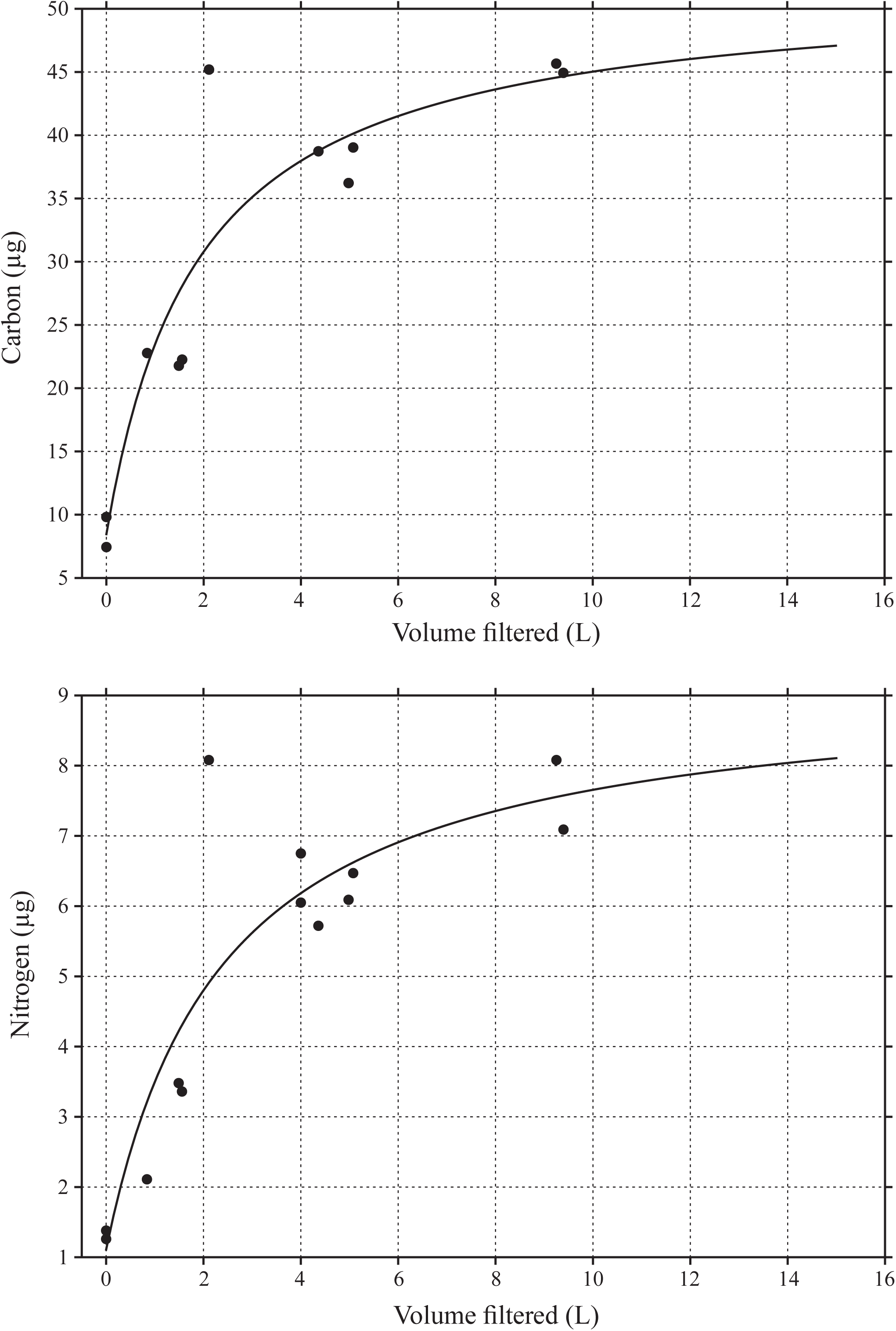
Filtered media blanks for particulate carbon and nitrogen measurements, as a function of volume of media filtered. Filtered media (47 mm diameter, 0.2 µm pore-size Nuclepore PC membrane followed by 0.22 µm pore-size, Sterivex-GP polyethersulfone membrane) was used as the source for procedural blanks filtered through combusted GF-75 glass-fiber filters to correct for adsorption of (A) dissolved organic carbon or (B) inorganic nitrogen. Carbon data fit to a rectangular hyperbolic saturation function: 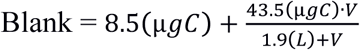 for medium volume (V) in liters. Nitrogen data fit to a rectangular hyperbolic saturation function: 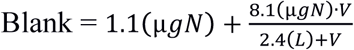 for medium volume (V) in liters.

**Figure S4.**
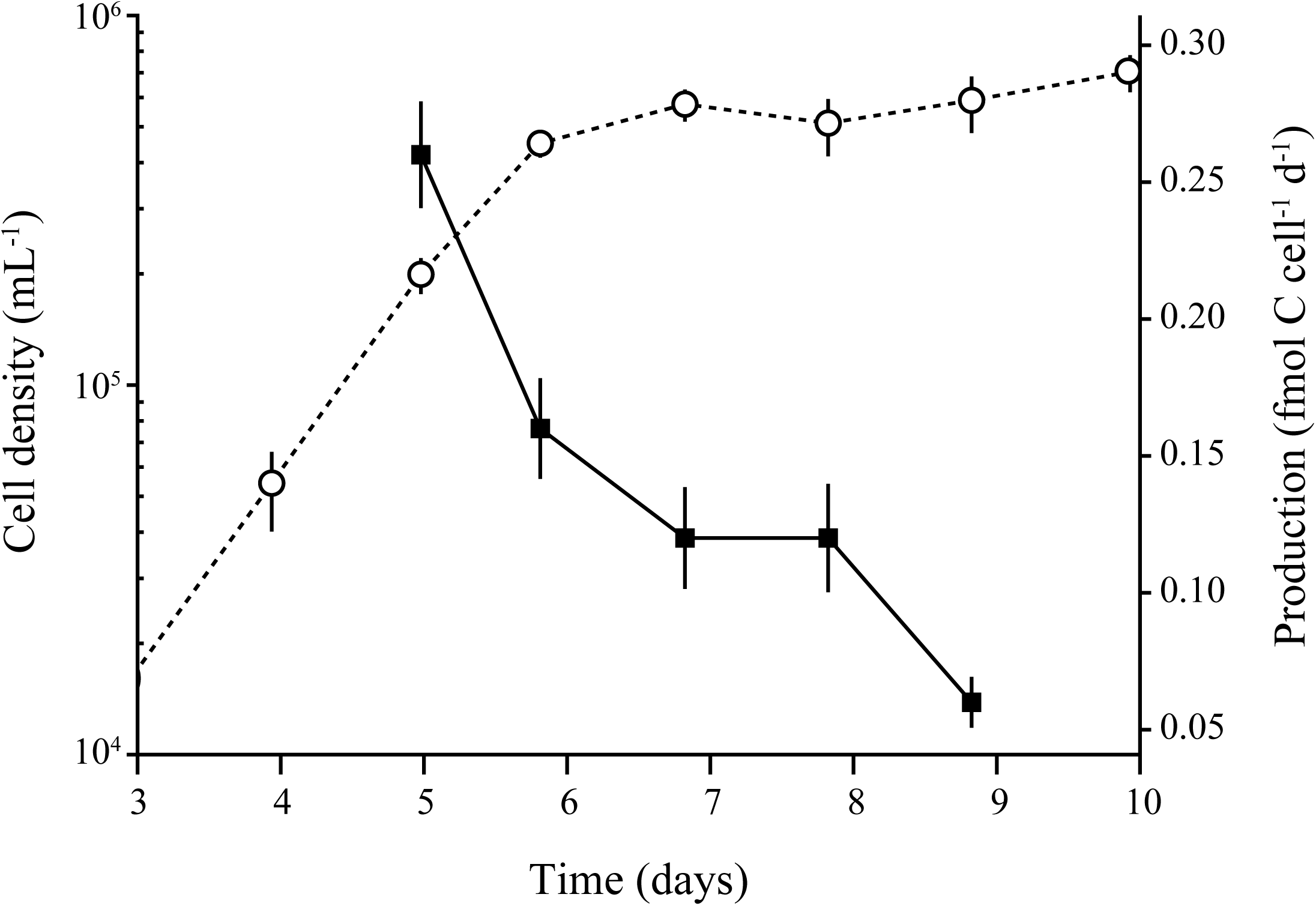
Bacterial production versus batch culture growth of strain HIMB114. Bacterial production (measured by ^3^H-Leu incorporation) during batch culture growth of a single 10 L culture of strain HIMB114, sampled throughout late exponential phase and into stationary phase. Open circles and dotted line indicate cell density, while filled squares and solid line indicate bacterial production.

**Figure S5.**
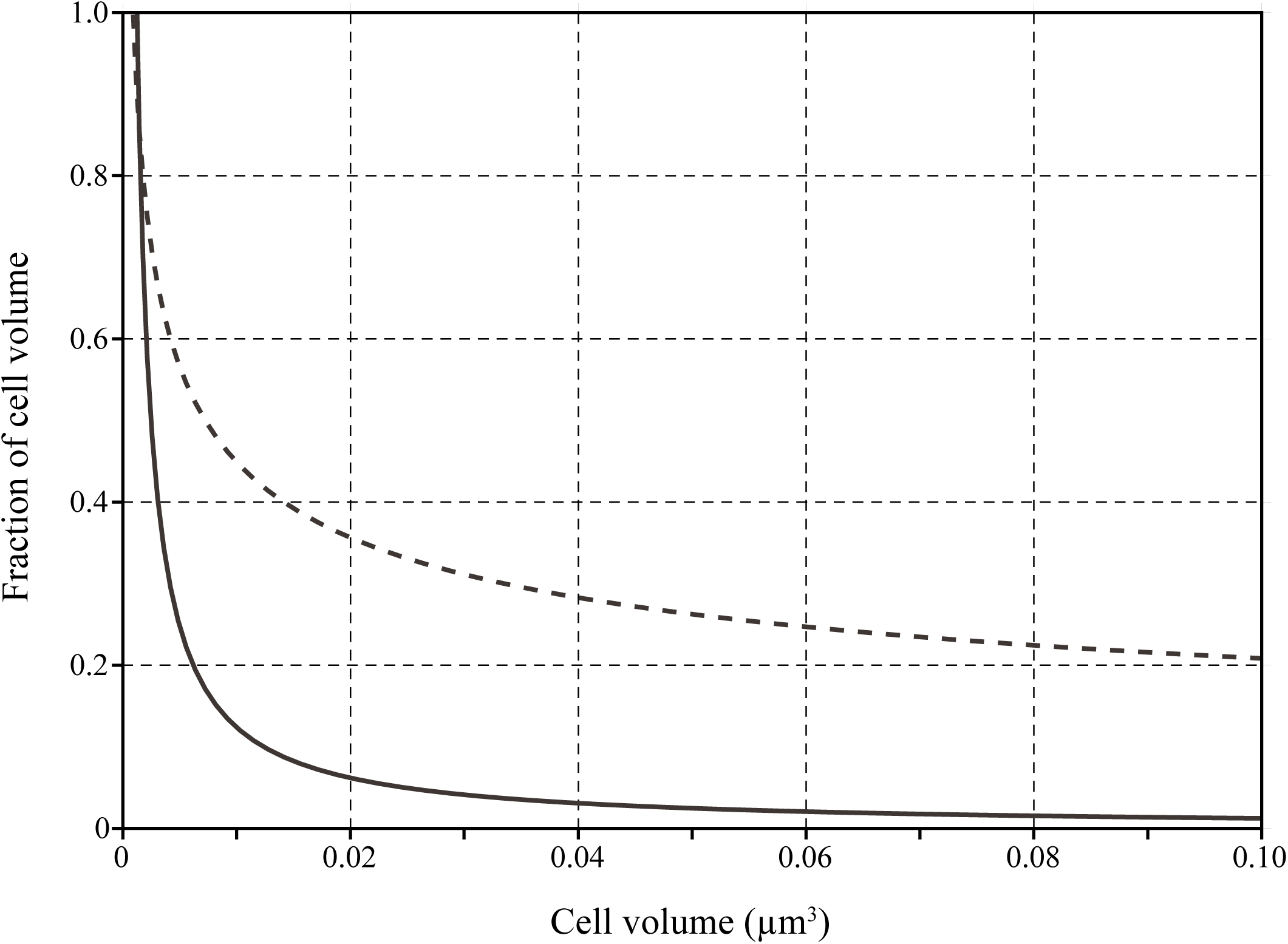
Volume scaling of the fraction of total cell volume occupied by the genome (solid line) and the cell envelope (dashed line) for a theoretical bacterial cell. Cell size parameters calculated assuming spherical cells, with volume of the cell envelope calculated assuming a spherical shell with an envelope thickness of 20 nm. Genome volume calculated assuming a genome length of 1.237 Mbp, and a DNA volume per base pair of 1 nm^3^. At the volume of the most frequent HIMB114 cell (0.05 µm^3^), the cell envelope and genome take up 26% and 2.5% of the cell volume, respectively.

**Figure S6.**
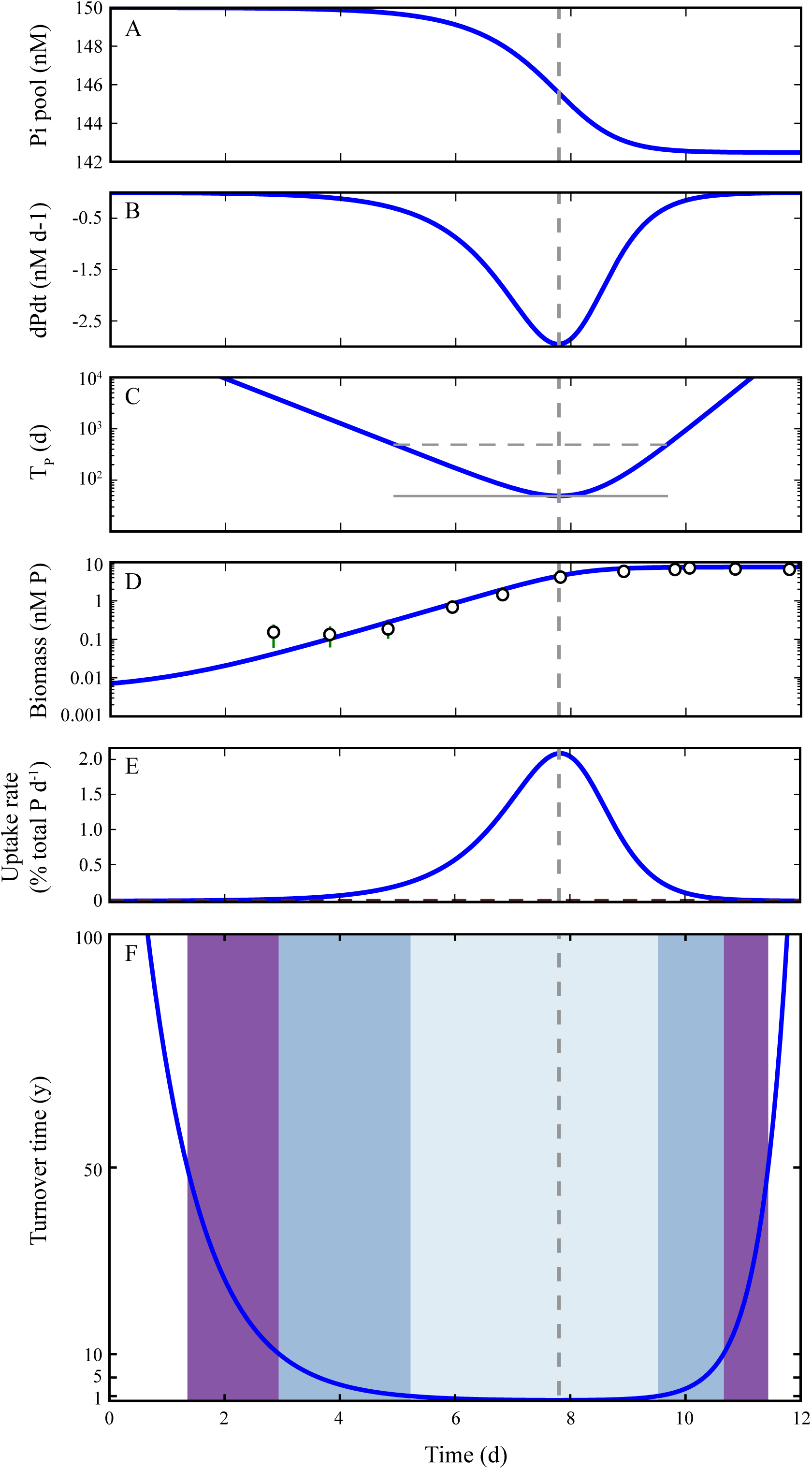
Batch growth and nutrient dynamics numerical model results (blue lines) for a batch culture of HIMB114, using phosphate as a sole source of phosphorus. (A) Phosphate (Pi) pool concentration; (B) rate of change of the Pi pool 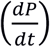 due to uptake by cells; (C) Pi pool turnover time (T_P_), with gray horizontal solid line indicating the minimum turnover time of 50 d, and gray dashed horizontal line showing when the turnover time is one order of magnitude above the minimum (500 d); (D) culture biomass as phosphorus concentration, with actual P-biomass calculated from cell counts from the culture (circles); (E) phosphate uptake rate dynamics, assuming phosphate as a sole source of phosphorus. The observed uptake rate value is plotted as a horizontal dashed gray line near the bottom of the y-axis, measured late on day 7 at 0.004% total P d^−1^. This intersects the modeled rate on day 11.6, nearly four days following the observed rate. (F) Similar to panel C, but with Pi pool turnover time in units of years and on a linear scale. Colored areas indicate time windows for turnover times less than 1 (light blue), 10 (blue), and 50 (purple) years. For all panels, the gray dashed vertical line indicates the time of highest uptake rate of phosphate, or equivalently the shortest turnover time. This corresponds closely to the actual timing of the measured Pi uptake experiments from the HIMB114 batch cultures, shown by the cell density circle closest to the vertical line in panel D.

**Figure S7.**
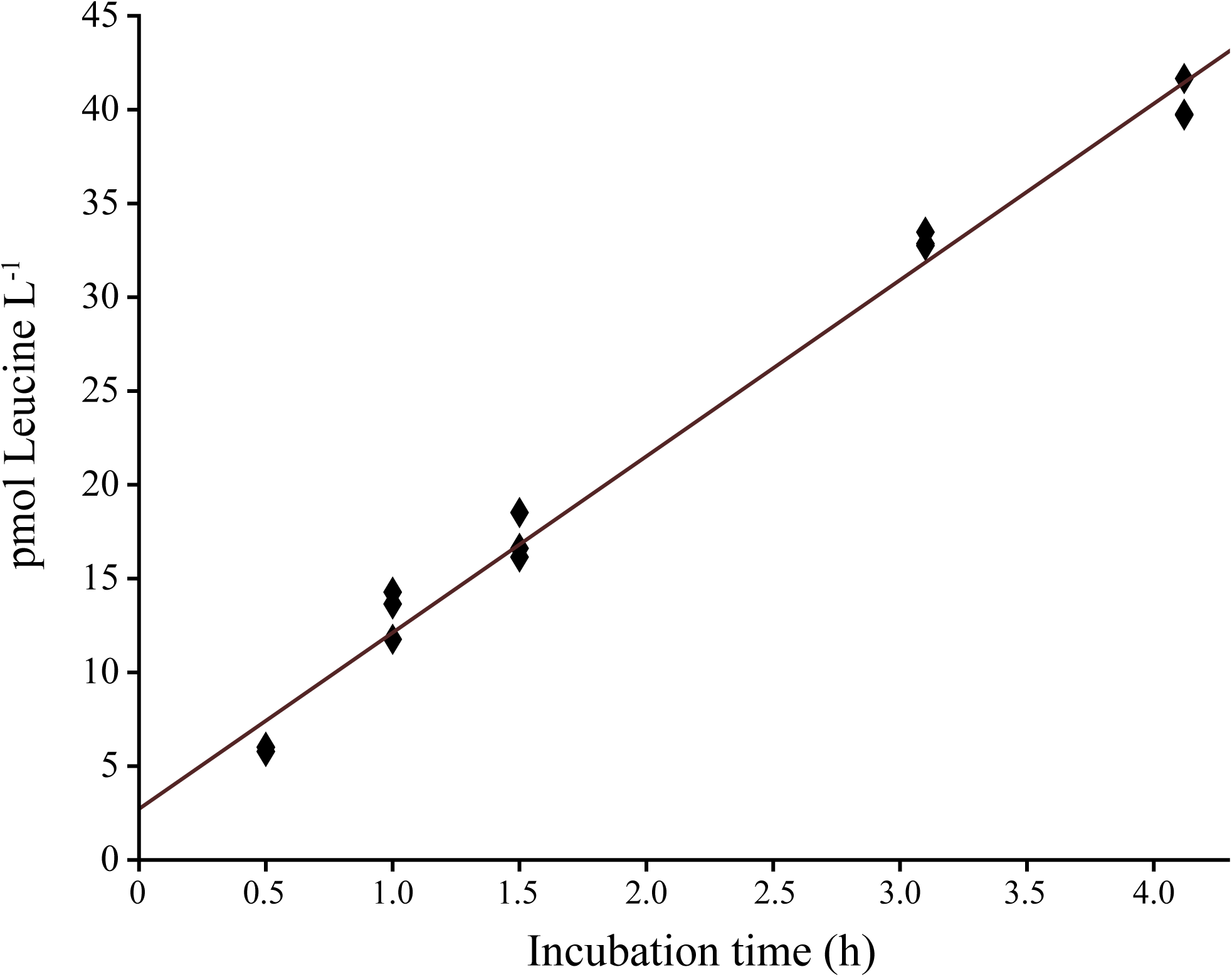
^3^H-Leucine incorporation by HIMB114 as a function of incubation time.

#### Cell enumeration and image analysis

Cell biovolume (V) was calculated from cell length (L) and width (W) using a bulk geometric function for a prolate spheroid, 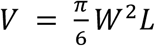. Although the bulk geometric function for a prolate spheroid has been shown to overestimate volumes for objects with similar shape morphology to HIMB114 by about 50%, it is likely the most appropriate bulk formula to use, and is better than using an equation for a cylinder with hemispherical caps, which can overestimate volumes for these shapes by 100% (Sieracki, Viles, & Webb, 1989).

#### Phosphate uptake

Control tests were made for the non-biological adsorption of ^33^P-tracer to both Nuclepore PC and Supor PES membranes (Pall Corp.) with and without the high-phosphate pre-saturation step. PC membranes were found to be superior to PES membranes for lowering control background, retaining just 0.0001% and 0.05% of the total ^33^P activity with and without the high-phosphate pre-saturation step, respectively. The Supor PES membranes retained significantly more of the ^33^P-phosphate radiotracer, 0.07% and 0.34% of the total activity with and without the high-phosphate step, and with much higher variance between replicates than for the PC membranes. Therefore, PC membranes were chosen as the preferred filters for reducing both the background and sample variance for ^33^P-phosphate uptake measurements.

#### Batch growth and nutrient dynamics model

A numerical model to describe the dynamics of batch culture growth was constructed to provide insights and theoretical comparisons for the measured PO_4_^3-^ uptake rates and turnover times under batch growth conditions. The model is cast in units of P concentration, in order to describe the uptake of PO_4_^3-^ and growth of the culture with the assumption that PO_4_^3-^ is the sole source of P for the culture.

The prognostic equation for the culture cell density (*x*) is described by a second order logistic equation:

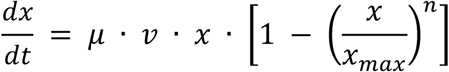

where *n* is the exponential constant for logistic growth, which we set to 2 after investigating the growth curves for n = 1 to 4 (an exponent higher than 1 is needed to correctly model the observed, rapid transition from exponential to stationary phase); *x*_*max*_ is the observed maximum cell density for the culture; *v* is the rectangular hyperbolic Monod function for phosphate limited growth, which never came into effect here because the phosphate pool never approached, within an order of magnitude, the assumed phosphate half-saturation constant for growth of *Kµ* = 1 nM P. The growth rate function, *µ*, is used to describe the transition from the lag phase to the exponential maximum growth rate phase, and is also described by a logistic equation:

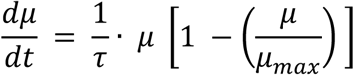

where *τ* is the timescale for the lag phase transition to exponential growth (1 d), and *µ*_*max*_ is the observed exponential phase growth rate. To convert from cell density to P-based biomass units, the measured P cell quota (*Q*_*P*_) is assumed to be constant throughout the growth curve, and the growth of cell biomass is directly coupled to the uptake of P from the phosphate pool:

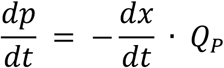

with observed initial concentrations of *P* and *x* used to initialize the model. The model was stepped forward with a time step of 30 minutes for 12 days using a simple Newton numerical scheme.

Because of the inherent non-linear dynamics of batch growth conditions, the theoretical nutrient uptake rates and turnover time may be quite dynamic, potentially changing by an order of magnitude on a daily timescale, making the interpretation of a measured rate on any particular day difficult. Results from the model matched reasonably well with the observed growth curve, and also confirmed that the time of maximum PO_4_^3-^ uptake rates (Fig. S6E) and minimum turnover time (Fig. S6C) occur at the end of the exponential phase of growth (Fig. S6D). This corresponded quite closely to when the PO_4_^3-^ uptake measurements were measured on this culture (Fig. S6D). The maximum theoretical specific uptake rate calculated by the model was 2×10^−2^ d^−1^, or 3 nmol P L^−1^ d^−1^, compared to the observed specific rate of 4×10^−5^ d^−1^, or 6 pM P L^−1^ d^−1^, measured very close to this time. The theoretical uptake rate falls off exponentially on either side of the maximum (Fig. S6E). The model also indicates that the minimum turnover time for the PO_4_^3-^ pool, occurring on day 7.8, should be close to 50 days, again increasing exponentially around the minimum. The actual turnover time measured at the end of day 7 was 70 years (range from 50 to 150), about 500 times the minimum turnover time. Even if the lower estimate of 50 years is used, the timing would need to be off by over 3.5 days to measure a turnover time of that scale (Fig. S6).

Knowing what range of PO_4_^3-^ turnover times to expect on a theoretical basis is quite difficult in batch cultures because of the non-linear dynamics, for which there is no clearly superior choice of functional parameterization for modeling batch culture growth (Zwietering et al., 1990). This creates a large uncertainty, of likely an order of magnitude, in the modeled uptake rates, which are particularly sensitive to the timing within the growth curve. Nevertheless, the turnover time was measured at what should be quite close to the time in the growth curve that would coincide with the minimum turnover time for PO_4_^3-^, and yet the measured turnover times were at least 500 times larger than expected.

#### Bacterial production

Typically, quadruplicate 1.5 mL sample volumes and duplicate blanks were added into 2 mL microcentrifuge tubes, followed by 2 µL of ^3^H-3,-4,-5-Leucine (106 Ci mmol^−1^; 5 mCi mL^−1^; PerkinElmer) to a final concentration of 60 nmol Leu L^−1^, mixed well, and incubated for 2.5 h under the same temperature and light conditions as the original cultures. Blanks were killed with trichloroacetic acid (TCA, 5% final) before the addition of ^3^H-Leu. Incubations were stopped by the addition of TCA (5% final, 4 °C), centrifuged (14,000 rpm at 4 °C for 15 min), rinsed with 1 mL 5% TCA (4 °C), centrifuged again (14,000 rpm at 4 °C for 5 min), then rinsed with 1 mL 80% ethanol (4 °C) and centrifuged (14,000 rpm at 4 °C for 5 min). Pellets were allowed to dry overnight at room temperature in a fume hood before adding 1 mL of scintillation cocktail (UltimaGold LLT; PerkinElmer), vortexed, and allowed to sit for at least four days before making final activity counts (PerkinElmer Tri-Carb Liquid Scintillation counter), as the activity was observed to increase over the first two days after adding cocktail. Activity counts were converted to leucine concentration based on the specific activity of the isotope and calibrated to a ^3^H standard. Prior to using single 2.5-hr time point incubations, the linearity of ^3^H-Leu incorporation by the HIMB114 culture was tested over 4 h and found to be quite linear over that time period (Fig. S7).

#### Oxygen respiration

Oxygen concentration measurements were made based on the mass spectrometric determination of the ratio of oxygen to argon. Briefly, the seawater sample was continuously pumped across a permeable membrane under vacuum, allowing dissolved gasses to diffuse across the membrane that were detected by an in-line mass spectrometer. An equilibrated seawater standard was used for calibration, and the oxygen concentration was calculated by the change in the O_2_/Ar ratio referenced to the initial, time zero, oxygen concentration. Oxygen respiration rates were then calculated by linear least squares regression of oxygen concentration versus incubation time over two-day periods.

#### Cellular Elemental Analysis

**Carbon and Nitrogen.** Preliminary tests of filtration methods indicated that filtration by even the lowest of vacuum pressure retained undetectable cells, while filtration by gravity retained at most 50% of cells. This was slightly better than very slow (5 mL min^−1^) peristaltic pump filtration, which was comparable to the rate of filtration by gravity. Because of the very slow filtration rates by gravity in combination with the large volumes that needed to be filtered, filtrations were conducted in a 4 °C walk-in cold room in order to stop cellular metabolism, and carried out over a period of 8 days of continuous gravity filtration. Pump speeds were adjusted to keep pace with the gravity filtration rates, which started at 3.5 mL min^−1^ and gradually slowed to 2 mL min^−1^ by day 5. Once filtration rates slowed to below 1 mL min^−1^ (day 8) the filtration was stopped.

For procedural blanks, batch cultures were 0.2 µm-filtered twice to remove all cells (0.2 µm pore-size, 47 mm-diameter Nuclepore PC membrane followed by 0.22 µm pore-size, Sterivex-GP PES membrane) and the filtrate used to construct dissolved carbon and nitrogen blank saturation curves by filtering 0, 1, 2, 5, and 10 L of sterile-filtered media through combusted GF-75 filters. These filters were then analyzed for carbon and nitrogen along with the sample filters (Fig. S3).

**Phosphorus.** Tubes containing sample filters were combusted for 9 h at 450 °C in order to convert organically bound phosphorus to inorganic phosphate, which was then extracted in 10 mL of 0.15 M HCl for 1 h at room temperature. Sub-samples (5 mL) of the acid extract were reacted with 0.5 mL of a molybdate mixed reagent solution to develop the blue phospho-molybdate complex. The molybdate mixed reagent was made by combining 10 mL of 30 g L^−1^ ammonium paramolybdate solution, 25 mL of 5N sulfuric acid, 10 mL of 5.4 wt.% ascorbic acid solution, and 5 mL of 1.7 g L^−1^ potassium antimony-tartrate solution. After allowing one h for color development, absorbance was measured on a spectrophotometer at 880 nm using a 1 cm path length quartz cell. Phosphorus concentrations were calculated using a phosphate standard curve, with the mean absorbance of the media procedural blanks subtracted from the sample absorbance.

